# MemBrain v2: an end-to-end tool for the analysis of membranes in cryo-electron tomography

**DOI:** 10.1101/2024.01.05.574336

**Authors:** Lorenz Lamm, Simon Zufferey, Hanyi Zhang, Ricardo D. Righetto, Florent Waltz, Wojciech Wietrzynski, Kevin A. Yamauchi, Alister Burt, Ye Liu, Antonio Martinez-Sanchez, Sebastian Ziegler, Fabian Isensee, Julia A. Schnabel, Benjamin D. Engel, Tingying Peng

**Affiliations:** Helmholtz Munich, German Research Center for Environment and Health, Munich, Germany; Biozentrum, University of Basel, Basel, Switzerland; School of Computation, Information and Technology, Technical University of Munich, Munich, Germany; Department of Biosystems Science and Engineering, ETH Zürich, Basel, Switzerland; Swiss Institute of Bioinformatics, Basel, Switzerland; MRC Laboratory of Molecular Biology, Cambridge Biomedical Campus, Cambridge, UK; Department of Structural Biology, Genentech, South San Francisco, CA, USA; Department of Information and Communications Engineering, Faculty of Computers Sciences, University of Murcia, Murcia, Spain; German Cancer Research Center (DKFZ), Division of Medical Image Computing, Heidelberg, Germany; Helmholtz Imaging, German Cancer Research Center (DKFZ), Heidelberg, Germany; School of Biomedical Engineering and Imaging Sciences, King’s College London, London, UK

**Keywords:** cryo-electron tomography, membrane segmentation, particle localization, deep learning, spatial analysis, MemBrain

## Abstract

Cryo-electron tomography (cryo-ET) provides unique insights into macromolecular complexes in their native environments, yet membrane analysis remains a major bottleneck due to low signal-to-noise ratios, missing wedge artifacts, and the complexity of membrane-associated proteins. Existing tools often require extensive manual annotation, struggle with generalization across datasets, and lack integrated solutions for segmentation, protein localization, and quantitative analysis. We introduce MemBrain v2, a deep learning-enabled framework that unifies these tasks into a streamlined pipeline. MemBrain-seg leverages a diverse, collaboratively generated training dataset and specialized model training strategies to achieve generalizable membrane segmentation across variable tomographic conditions. MemBrain-pick enables data-efficient localization of membrane-bound proteins by integrating geometric constraints with deep learning, reducing the need for extensive manual annotation. MemBrain-stats provides quantitative insights into protein distributions, computing spatial metrics to analyze intra-membrane particle organization. MemBrain v2 integrates seamlessly into cryo-ET workflows, providing an accessible and structured approach to membrane analysis. The full package is available at https://github.com/CellArchLab/MemBrain-v2.

## Introduction

Cryo-electron tomography (cryo-ET) is a powerful technique for imaging the molecular environment inside native cells in three dimensions (3D) at sub-nanometer resolution^1^. By capturing all densities inside a cellular volume frozen in vitreous ice, cryo-ET provides detailed insights into the structures, interactions, and spatial arrangements of diverse organellar and macromolecular components. However, the biological complexity captured by cryo-ET also presents significant challenges for annotation and analysis^2^. Membranes and their embedded protein complexes play fundamental roles in numerous cellular processes, including the ER-Golgi secretory pathway^3–5^, autophagy^6,7^, vesicular transport^8^, synaptic transmission^9–12^, energy conversion in mitochondrial cristae and chloroplast thylakoids^13–22^, organelle contact sites^23,24^, cell membrane protrusions^25,26^, viral replication^27–33^, and bacterial cell division^34,35^. Accurate segmentation of membranes and precise localization of membrane-associated proteins are essential for understanding how membrane architecture and molecular organization drive cellular functions.

Membrane analysis in cryo-ET is challenging due to the inherently low signal-to-noise ratio (SNR), which obscures structural details and complicates accurate interpretation. Additionally, the missing wedge effect – caused by limited angular sampling – introduces anisotropic distortions that particularly affect membrane structures oriented perpendicular to the electron beam, making them difficult to resolve. While computational approaches for denoising^36–39^ and missing wedge correction^40,41^ attempt to mitigate these issues, their restorations are not always reliable and may introduce artifacts.

Several methods attempt to tackle membrane segmentation in cryo-ET, yet each comes with inherent limitations. Classically, TomoSegMemTV^42^ incorporates local membrane curvature via tensor voting, but can struggle in areas with complex or rapidly varying membrane shapes. Deep learning-based approaches, particularly U-Net-based^43–46^ models, have driven significant advancements. However, many implementations lack generalizability, because they have been trained on only a single cell or membrane type. Therefore, specialized models are often trained for single projects or datasets^47^. TARDIS^48^ represents a promising step toward more generalizable segmentation by incorporating diverse datasets into training and offering pretrained models, making segmentation tools more accessible. Despite these advancements, it remains an open challenge to develop a widely applicable approach that ensures robust segmentation across diverse membrane architectures, tomographic conditions, and biological contexts.

Beyond membrane segmentation, the automated localization of membrane-associated particles (e.g., integral membrane proteins) in cryo-ET remains a major challenge. Because these particles are embedded in the lipid bilayer, they are difficult to distinguish from the surrounding membrane. Classical template-matching approaches^49–51^ often fail in this context, as strong membrane contrast dominates cross-correlation scores, reducing detection accuracy. Deep learning methods offer an alternative, with several CNN-based approaches recently introduced for particle localization in cryo-ET^52–57^. However, many of these heavily rely on extensively annotated volumes, which are labor-intensive and often not technically feasible to generate. These models also struggle with sparsely annotated membranes, limiting their generalizability. The high variability in particle appearances further complicates training of generalist models, making each individual membrane annotation particularly valuable and motivating the need for specialized models that can operate with individual membrane annotations. To streamline this annotation process, MPicker^58^ flattens membranes into 2D stacks, but this transformation can introduce distortions and fail to preserve physical distances. To address these issues, annotation tools such as membranorama^17,59^ and Surforama^60^ project tomographic densities onto 3D meshes, allowing for more interactive exploration and particle localization. Among existing tools, MemBrain v1^61^ was designed to leverage these limited annotations, but its performance remains constrained by its small receptive field. A more refined, interactive approach could better leverage sparse annotations to improve both the accuracy and efficiency of particle localization.

Quantitative tools for analysis of membrane segmentations include PyCurv^62^, which estimates membrane curvature, and a surface morphometrics pipeline^13^, which extracts properties such as inter-membrane distances. PyOrg^63^ analyzes protein distributions but does not directly relate them to membrane features. Thus, these tools analyze either membrane geometry or protein distributions separately, lacking an integrated approach to quantitatively assess protein-membrane interactions.

To address the challenges described above, MemBrain v2 unifies cryo-ET membrane analysis in a single integrated framework: **MemBrain-seg** produces generalizable membrane segmentations using a collaboratively generated, diverse ground truth dataset. **MemBrain-pick** efficiently localizes membrane-bound particles with interactive annotation support via Napari and Surforama. **MemBrain-stats** computes descriptive statistics such as particle concentrations and geodesic nearest neighbor distances along the segmented membrane. Together, these modules enable end-to-end analysis of membranes from segmentation to particle localization and statistical analysis (Fig. 1). By combining versatility with usability, MemBrain v2 provides an intuitive solution that can be applied to diverse data sources to explore a broad spectrum of biological questions.

**Figure 1.**
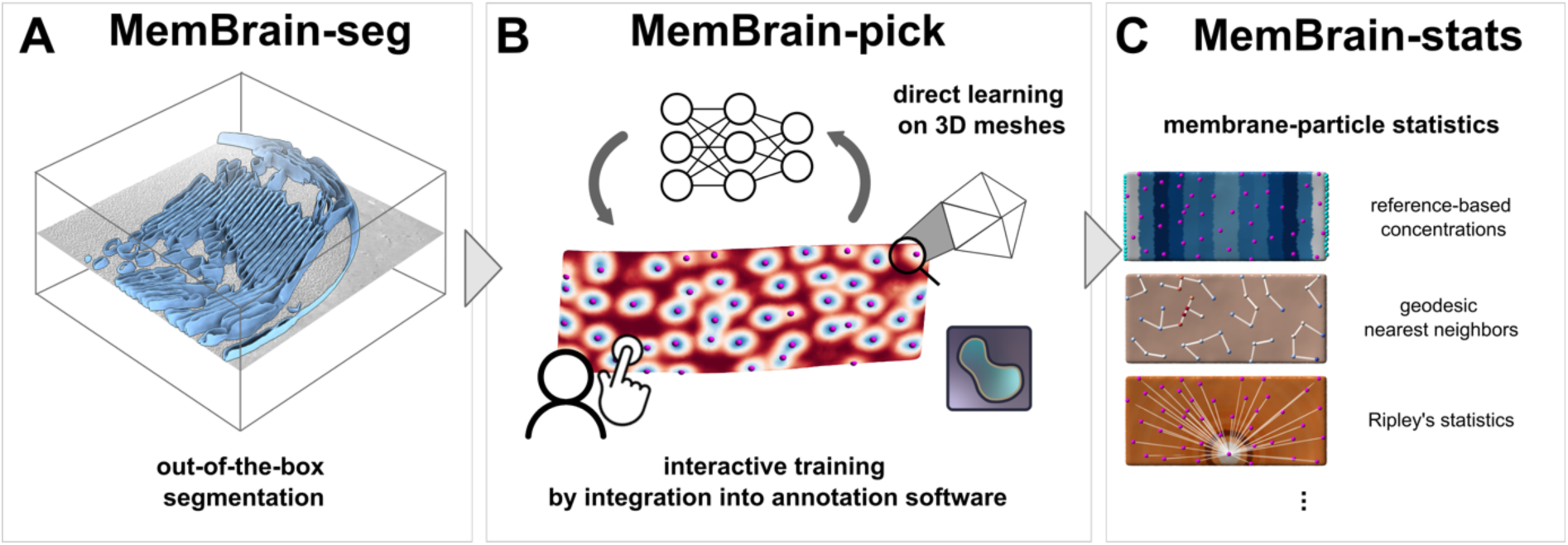
MemBrain v2 provides an end-to-end pipeline for analyzing membranes and membrane-associated particles in cryo-ET data. **A**: MemBrain-seg produces generalizable membrane segmentations out-of-the-box. **B**: MemBrain-pick efficiently localizes membrane-associated particles by iterative annotation and training directly on mesh surfaces. **C**: MemBrain-stats evaluates the outputs of MemBrain-seg and MemBrain-pick, providing particle-surface metrics.

## Results

### Overview of MemBrain v2 modules

MemBrain v2 is a modular pipeline designed to streamline the analysis of membranes and their associated proteins in cryo-ET datasets. Its three core modules — MemBrain-seg, MemBrain-pick, and MemBrain-stats — work together to enable membrane segmentation, particle localization, and quantification. MemBrain-seg (Fig. 1A) employs a U-Net-based approach to achieve robust membrane segmentation across a variety of experimental and tomography setups. It is trained on a diverse, iteratively refined dataset, which was generated using careful manual annotations and corrections in close collaboration with the community to ensure broad coverage of different membrane appearances. Together with our cryo-ET-specific data augmentations and membrane-focused loss functions in the network training, this approach ensures accurate and continuous membrane delineation, facilitating visualization and downstream analysis. MemBrain-pick (Fig. 1B) specializes in the efficient localization of membrane-bound particles (e.g., membrane proteins). By training a neural network to operate directly on membrane surfaces, it incorporates the membrane geometry into its prediction and thus reduces the search space, enhancing both accuracy and data efficiency. Its integration with interactive Napari^64^ tools, such as Surforama^60^, allows for rapid annotation and refinement of particle positions, facilitating seamless transitions between ground truth (GT) generation and model training. MemBrain-stats (Fig. 1C) leverages the outputs of MemBrain-seg and MemBrain-pick to provide quantitative insights into particle distributions on membranes. It computes key metrics such as particle concentrations, geodesic nearest-neighbor distances, and Ripley’s statistics. By linking membrane morphology with protein organization, MemBrain-stats enables investigation of the structural and functional relationships in cryo-ET datasets. MemBrain v2’s modular design ensures both flexibility and accessibility, featuring straightforward command-line interfaces and seamless integration with Napari plugins for interactive annotation and visualization.

### MemBrain-seg: a generalized approach for membrane segmentation

MemBrain-seg is a U-Net-based program that generates 3D membrane segmentations from input tomograms with a single command (Fig. 2A, Supp. Fig. S1). It delivers robust segmentation performance across diverse tomograms containing a variety of organelles from various species, acquired with different microscope types, imaging settings, and processing conditions, ranging from raw to denoised tomograms (Fig. 2E, Supp. Fig. S2). While strong segmentation performance on thylakoid membranes was expected due to the high representation of spinach chloroplasts in our training data (see Supp. Fig. S3, <u>Supp.</u> Table 1), MemBrain-seg also achieves satisfactory results on vesicular structures and convoluted mitochondrial membranes. This adaptability is further demonstrated in the CZII data portal^65^ (Supp. Fig. S4), where MemBrain-seg was applied to all tomograms present in the portal, allowing users to conveniently inspect segmentations together with the corresponding tomograms in the browser.

**Figure 2.**
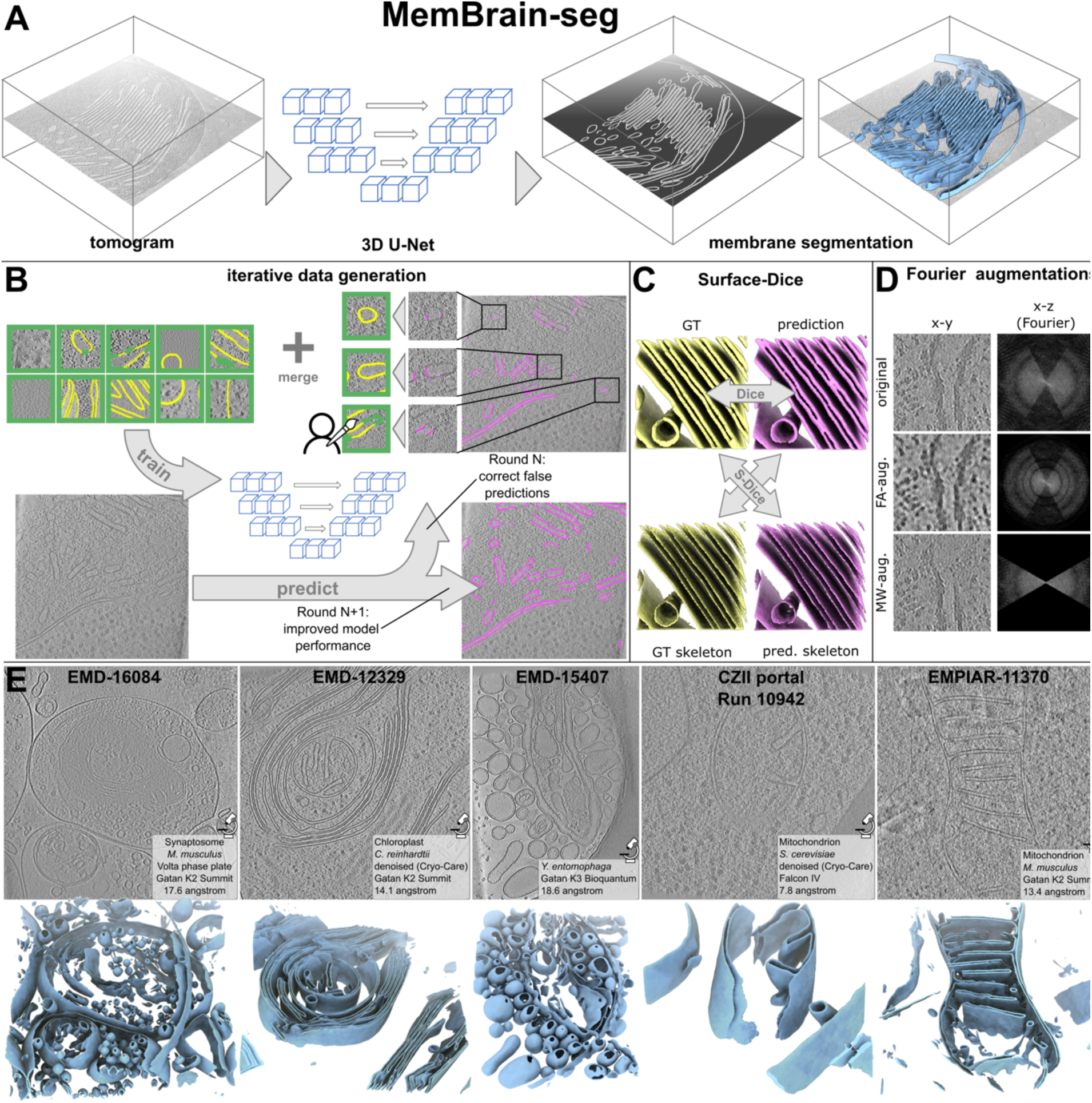
MemBrain-seg achieves accurate and generalizable membrane segmentation across diverse cryo-ET datasets through iterative active learning, membrane-specific loss function and Fourier-based augmentation. **A:** MemBrain-seg is based on a U-Net architecture and produces accurate 3D segmentations from input tomograms. **B**: Our diverse training dataset was generated iteratively via an active learning approach: For a model in training round N, we extracted patches in difficult regions, manually corrected the network prediction using membrane, background, and ignore labels (see Supp. Fig. S3), and merged them into our training dataset. The next round (N+1) shows improved segmentation performance. **C**: Surface-Dice evaluates segmentation quality by comparing the skeletons (see Supp. Fig. S15) of the predicted segmentation with the ground truth (GT) segmentation (yellow), and the GT skeleton with the predicted segmentation (pink). This emphasizes membrane continuity more efficiently than the conventional Dice metric, which directly compares voxel-wise overlap. **D:** MemBrain-seg’s Fourier-based augmentations: Fourier amplitude (FA) augmentation randomly rescales intensities of Fourier frequency bands, altering image contrast. The missing wedge (MW) augmentation randomly applies an artificial missing wedge in Fourier space to imitate the effects of the real missing wedge. **E**: Examples of MemBrain-seg predictions on diverse public datasets, demonstrating robust out-of-the-box segmentation performance across a wide range of membrane architectures in tomograms acquired with diverse instrumentation. Top row: slices through tomograms, bottom row: corresponding MemBrain-seg predictions (light blue). For more examples, see Supp. Fig. S2.

To ensure consistent and robust model performance, MemBrain-seg was trained on a diverse dataset, iteratively refined through manual corrections (Supp. Fig. S3). In each training round, segmentation predictions were reviewed and corrected to improve the dataset for subsequent iterations (<u>see</u> Fig. 2B). The first two rounds (1 and 2) focused on training patches from our *Spinacia oleracea* (“Spinach”, EMPIAR-12612) and *Chlamydomonas reinhardtii* (“Chlamy”, EMPIAR-11830) datasets, followed by contributions from external collaborators (“Collaborators”) to introduce additional diversity. To further enhance robustness, synthetic data from publicly available generators^39,66^ (“Synthetic”) and patches from the DeePiCt dataset^44^ (“DeePiCt”, EMPIAR-10988) were integrated in the last two training rounds (4 and 5, respectively). Expanding the dataset incrementally across different tomography sources further enhanced the model’s generalization, as demonstrated by its progressively improving performance across datasets with each additional training round (Supp. Fig. S5). Here, we monitored performance by calculating the Dice score (a standard metric for segmentation accuracy) for different test datasets. The improved generalization is particularly evident in the “DeePiCt” test dataset, where Dice scores improved from 39% to 66% (and up to 55% in rounds without any DeePiCt training data).

In addition to evaluating network performance across different training datasets, we also compared MemBrain-seg to TARDIS^48^, a recent method designed for membrane and filament segmentation. In our test datasets, MemBrain-seg achieved higher Dice scores compared to TARDIS (Supp. Fig. S5A). However, this comparison should be interpreted carefully because our GT annotations were generated by iteratively predicting with MemBrain-seg and manually correcting these predictions. As a result, a substantial portion of the GT voxels still originates directly from MemBrain-seg outputs, and may inherently favor its performance. To address this issue, we additionally assessed performance using fully synthetic datasets with absolute GT annotations, providing a more unbiased, but also less realistic, benchmark. Here, both MemBrain-seg and TARDIS achieved good values (66% in Round 3 vs. 59% Dice, respectively), hinting at strong generalization capabilities for both methods. To further mitigate potential bias — particularly differences in membrane thickness annotations, which may favor MemBrain-seg’s predicted segmentation thickness — we introduced the Surface-Dice score (Fig. 2C). Unlike standard Dice scores, which compare segmentations at the voxel level, Surface-Dice evaluates the structural consistency of predicted and GT segmentations by comparing their skeletons (see Supp. Fig. S6). This metric better captures membrane continuity and is invariant to annotation style (e.g., Supp. Fig. S7). Evaluations with Surface-Dice were consistent with the standard Dice assessments (Supp. Fig. S5A). By combining Dice and Surface-Dice metrics with our diverse dataset, we provide a publicly available benchmarking resource for membrane segmentation tools.

In addition to our base model, we provide versions trained with enhanced augmentations. These Fourier-based augmentations (Fig. 2D, Supp. Fig. S8B,C) simulate tomographic style variations and missing wedge distortions, improving model robustness. Specifically, our missing wedge augmentation artificially introduces an additional missing wedge in training patches to mimic real-world artifacts, while Fourier amplitude augmentation randomly rescales Fourier frequency bands to simulate tomographic variability. These enhancements improved generalization across diverse tomographic conditions (Supp. Fig. S5B). Notably, for round 4 data (i.e., excluding DeePiCt training data), we evaluated models trained with either Fourier amplitude or missing wedge augmentation. Fourier amplitude increased Surface-Dice scores on the DeePiCt test set from 55% to 59%, while missing wedge augmentation improved scores to 57%, demonstrating their effectiveness in adapting to unseen datasets. Overall, MemBrain-seg delivers strong out-of-the-box segmentation performance across a wide range of cryo-ET datasets. However, its performance may degrade when applied to datasets with substantial domain shifts, particularly under varying microscopy conditions. In such cases, fine-tuning the model on a specific dataset (e.g., with patches generated as shown in Supp. Fig. S3A) can further optimize segmentation quality. We demonstrate this capability in Supp. Fig. S9, where fine-tuning with only DeePiCt dataset patches in the training set improved Surface-Dice scores from 58% to 67%. To prevent overfitting, we continuously monitored performance on the full validation set depicting a broader range of membranes than the test set, using both the Dice score and Surface-Dice score. Notably, monitoring Surface-Dice during fine-tuning outperformed monitoring only Dice scores (67% vs. 60%, respectively), avoiding premature early stopping due to the different membrane thicknesses in the fine-tuning dataset (<u>Supp. Fig.</u> <u>S7</u>) and highlighting the effective adaptation of MemBrain-seg to domain shifts.

### MemBrain-pick: an interactive tool to efficiently localize membrane-associated particles

Identification of membrane-associated particles (e.g., integral membrane proteins or membrane-bound ribosomes) in cryo-ET data is a challenging task. In MemBrain-pick, we aim to facilitate this process by enabling a smooth transition from MemBrain-seg outputs to the area of interest for particle localization and training an automated model. With our MemBrain Napari plugin and its integrated 3D lasso functionality, users can isolate single membrane instances from full-tomogram segmentations (Fig. 3A, Supp. Fig. S1B), which can subsequently be visualized and annotated in Surforama^60,64^ (Fig. 3B) for manual annotation of particle locations.

**Figure 3.**
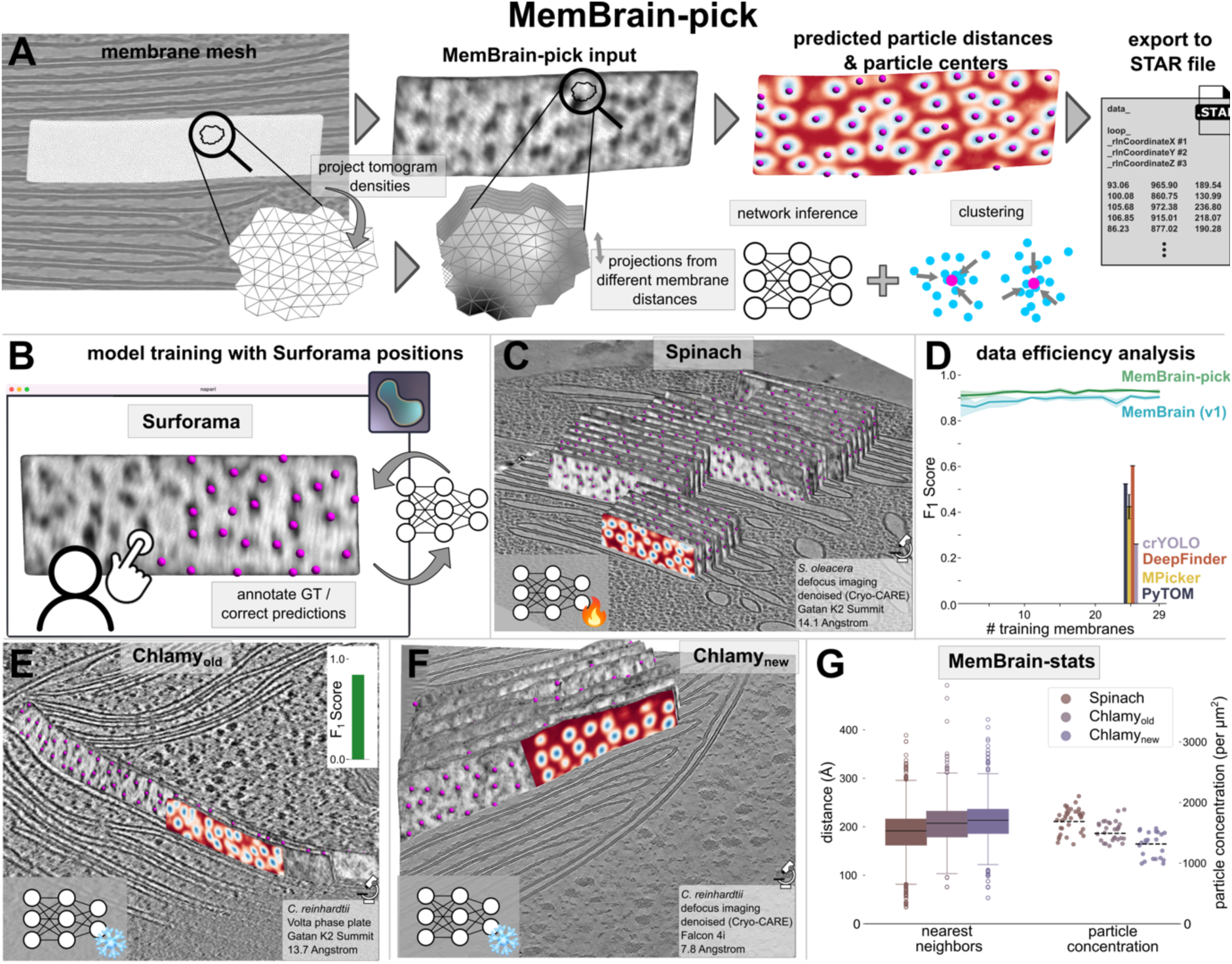
MemBrain-pick accurately localizes membrane-associated particles through efficient learning on mesh surfaces. **A:** MemBrain-pick projects tomographic densities onto a triangular mesh surface at multiple distances along the membrane normal vector. The DiffusionNet^64^-based architecture operates directly on the surface and predicts a heatmap depicting the distance to the nearest particle position (magenta). Particle positions are subsequently identified using Mean Shift clustering on the heatmap and exported as .star files. See detailed workflow in Supp. Fig. S10. **B:** Surforama interface enables interactive annotation and correction of MemBrain-pick outputs to generate or refine ground-truth particle positions. **C:** MemBrain-pick predictions on the Spinach test dataset after training with the Spinach training dataset. **D:** MemBrain-pick training results with incremental numbers of training membranes and comparison with other methods (see also Supp. Fig. S13). crYOLO and DeepFinder were only trained with 25 membranes (validation split based on tomogram), and MPicker with 29 membranes. Template matching was performed in PyTOM. **E:** Predictions on the Chlamyold dataset with the same model as in B. Inset shows the F1-Score performance evaluated with GT particle positions. **F:** Predictions on the Chlamynew dataset with the same model as in B. **G:** MemBrain-stats quantifies nearest neighbor distances and particle concentration (<u>Supp.</u> <u>Fig. S14</u>) for the predictions in B, C, and D.

These annotations allow the training of a specialized particle localization model: MemBrain-pick enables efficient and accurate localization of membrane-bound particles by integrating membrane geometry into the particle detection pipeline. It employs a DiffusionNet-based^67^ neural network to directly operate on membrane meshes with projected tomographic densities (Fig. 3A, Supp. Fig. S10), which reduces the search space to membrane-associated regions and improves detection efficiency. Additionally, this network design is aligned with the visualizations in Surforama, making the output more interpretable. The network predicts a heatmap representing the distance to the nearest particle center, which is then further processed with our score-guided mean shift clustering^68^ to predict precise particle positions (see example in Supp. Fig. S11).

We tested MemBrain-pick on stacked Spinach thylakoid membranes, demonstrating exceptional performance for Photosystem II (PSII) localization (Fig. 3C, D). Even with only a single annotated membrane, our model achieved robust performance with an F1-score of 91%, highlighting its ability to operate with minimal training data. In contrast, other deep learning approaches such as DeepFinder and crYOLO require extensive volumetric annotations (<u>Supp.</u> <u>Fig. S12</u>) and struggle with sparsely annotated training data (DeepFinder 60%, crYOLO 26% F1-score when trained with 25 membranes, Supp. Fig. S13B,C). Similarly, specialized approaches like MPicker (combined with EPicker, Supp. Fig. S13A, 42% F1-score) and MemBrain v1 (90% F1-score) also fail to match MemBrain-pick’s performance (93% F1-score when trained on 29 membranes). Template matching (performed in PyTOM^49,50^) achieved an F1-score of 52%.

To evaluate MemBrain-pick’s generalizability across datasets, we applied one of these Spinach-trained models (i.e., trained with 29 membranes) to stacked thylakoid membranes in two Chlamydomonas datasets (Chlamyold and Chlamynew). Despite differences in species and imaging setups, MemBrain-pick maintained robust localization with high F1-scores (84%) on the Chlamyold dataset (Fig. 3E). Predictions on the Chlamynew dataset (which lacked GT annotations) also appeared clean and plausible (Fig. 3F).

### MemBrain-stats: quantitative analysis of particle distributions

MemBrain-stats complements MemBrain-seg and MemBrain-pick by providing quantitative analysis of particle distributions on membrane surfaces. The module computes particle surface concentrations, geodesic nearest-neighbor distances, and spatial distribution statistics such as the Ripley’s functions (see Supp. Fig. S14). These metrics allow users to quantitatively interpret membrane protein organization within cryo-ET data, enabling deeper biological insights into membrane-associated processes.

Using MemBrain-stats, we quantified the distribution of the above-predicted thylakoid membrane particles and compared to previously published values of total particles in stacked thylakoids (Fig. 3G). For Spinach, our results closely align with the published particle concentration^19^ (Spinach vs. published: 1688 vs. 1714 particles/µm^2^). For Chlamydomonas, our predicted values are slightly higher than the reported concentration^17^ (Chlamyold and Chlamynew vs. published: 1492 and 1316 vs. 1292 particles/µm^2^). Because the previous studies only measured PSII-PSII nearest neighbor distances and neglected some densities marked as “unknown”, our computed nearest neighbor distances between all particles (corresponding to PSII + “unknown”) are slightly shorter (Spinach vs. published: 18.8 vs. 21.2 nm; Chlamyold and Chlamynew vs. published: 20.7 and 21.2 vs. 24.4 nm).

In summary, this streamlined MemBrain v2 workflow achieved comparable results to previously published manual analysis of stacked thylakoids, but with greatly accelerated speed and throughput that opens the door to studies with more biological conditions and reproducibility. Below, we apply this pipeline to analyze the higher-order organization of additional types of membrane-bound particles – phycobilisomes and ribosomes.

### Test application: phycobilisome organization on thylakoid membranes

Phycobilisomes are large light harvesting antennae, found in cyanobacteria and some species of eukaryotic algae, that capture light and transfer the excitation energy to thylakoid membrane-embedded photosystems. Phycobilisomes from different species were previously shown by cryo-ET to assemble into rows^69,70^, and the function of this higher-order organization in photosynthesis remains an open question. Previous studies of phycobilisome organization in cryo-ET relied heavily on manual annotations, making a large-scale analysis extremely time-consuming^71,72^. By applying MemBrain v2 to a high-resolution cryo-ET dataset of red algae chloroplasts (EMD-31244^71^), we demonstrate how our approach can efficiently extract particle positions and spatial patterns of phycobilisome chains (Fig. 4).

**Figure 4.**
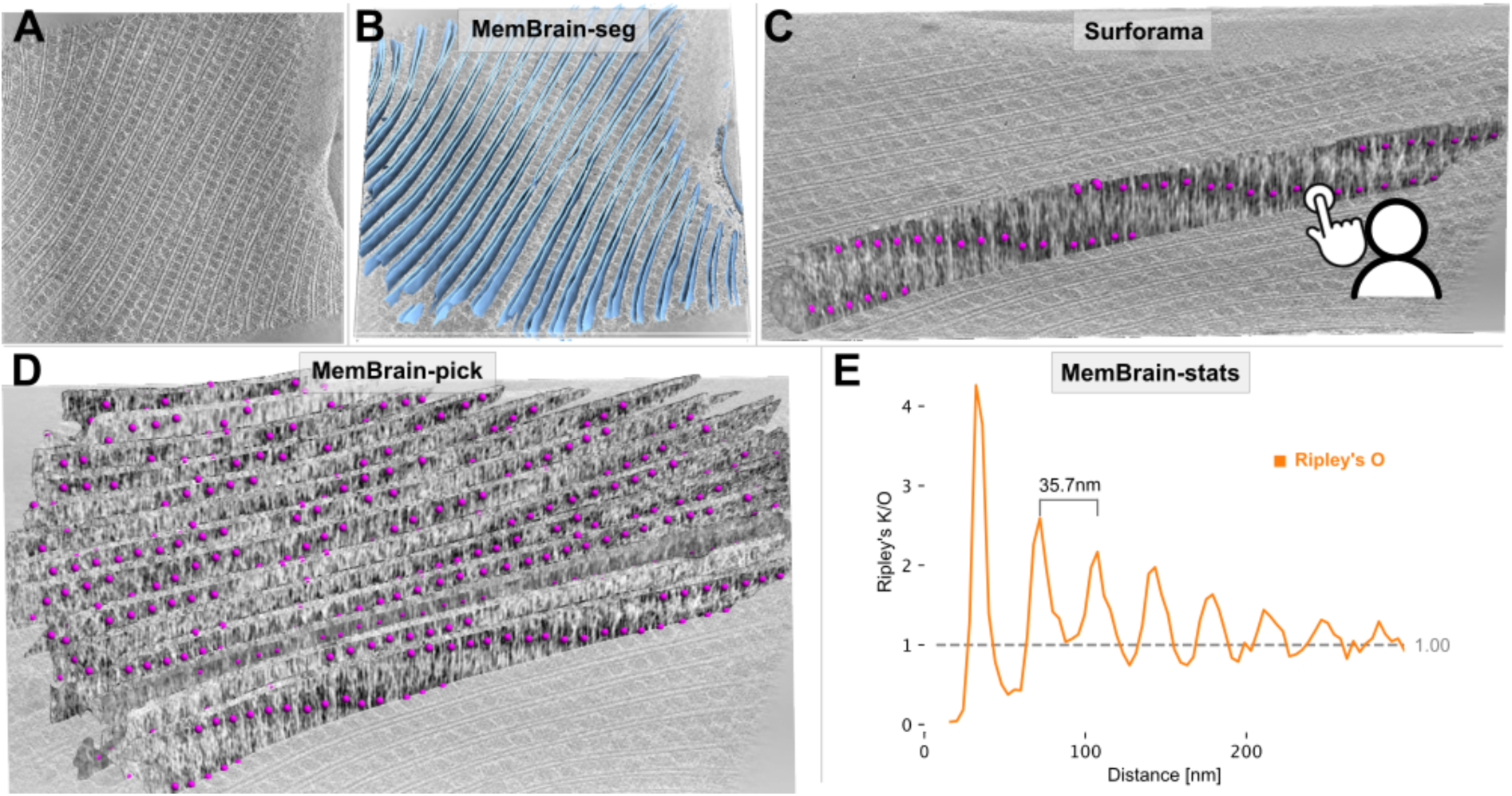
MemBrain v2 end-to-end workflow detects periodic phycobilisome organization. **A:** Raw tomogram slice of EMD-31244. **B:** Out-of-the-box MemBrain-seg segmentation (light blue). **C:** A single membrane instance can be visualized in Surforama and manually annotated with GT phycobilisome positions (magenta). **D**: MemBrain-pick localizes particles (trained with data from C) on all membranes in the tomogram. **E**: MemBrain-stats computes Ripley’s O statistic using the positions from D with a bin size of 5nm. The distance between peaks (35 nm) was measured to estimate chain unit spacings.

In a first processing step, MemBrain-seg consistently produced clearly separated, high-precision segmentations of all thylakoid membranes (Fig. 4B). For further processing, we utilized the MemBrain lasso tool (Supp. Fig. S1B) to perform connected component analysis, allowing us to extract individual membrane instances and visualize them in Surforama^60^. This interactive visualization enabled efficient manual annotation of phycobilisome chain unit positions in six selected membrane instances (Fig. 4C). These manually determined positions served as GT data to train a MemBrain-pick model. Once trained, MemBrain-pick was applied to the remaining 23 membranes in the tomogram, successfully identifying phycobilisome positions across the entire tomogram. The predicted positions exhibited a clear periodic arrangement of phycobilisomes along the membranes (Fig. 4D). To quantify this spatial organization, we applied MemBrain-stats to compute the Ripley’s O function, which confirmed the regular spacing of phycobilisomes (∼35 nm) (Fig. 4E). This periodicity is consistent with previous manually determined values (34.5 nm^71^), demonstrating the reliability of our automated approach. This end-to-end MemBrain analysis from raw tomograms to extracted phycobilisome chains can rapidly extract spatial information from large datasets to investigate native membrane organization in many cells and experimental conditions.

### Test application: poly-ribosome chain organization on nuclear envelope

In another end-to-end application of MemBrain v2, we analyzed ribosome distributions on the outer membrane of the nuclear envelope, aiming to identify patterns in their localization and orientation. The EMPIAR-11830^73^ dataset contains over 1800 tomograms capturing diverse organelles in *Chlamydomonas reinhardtii*. When analyzing tomograms containing the nuclear periphery, MemBrain-seg accurately segmented the nuclear envelope as well as all cytoplasmic membrane structures around it (Fig. 5A). Using the MemBrain lasso tool, we isolated the nuclear envelope membranes of interest and visualized them in Surforama, enabling efficient annotation of GT ribosome positions (Fig. 5B).

**Figure 5.**
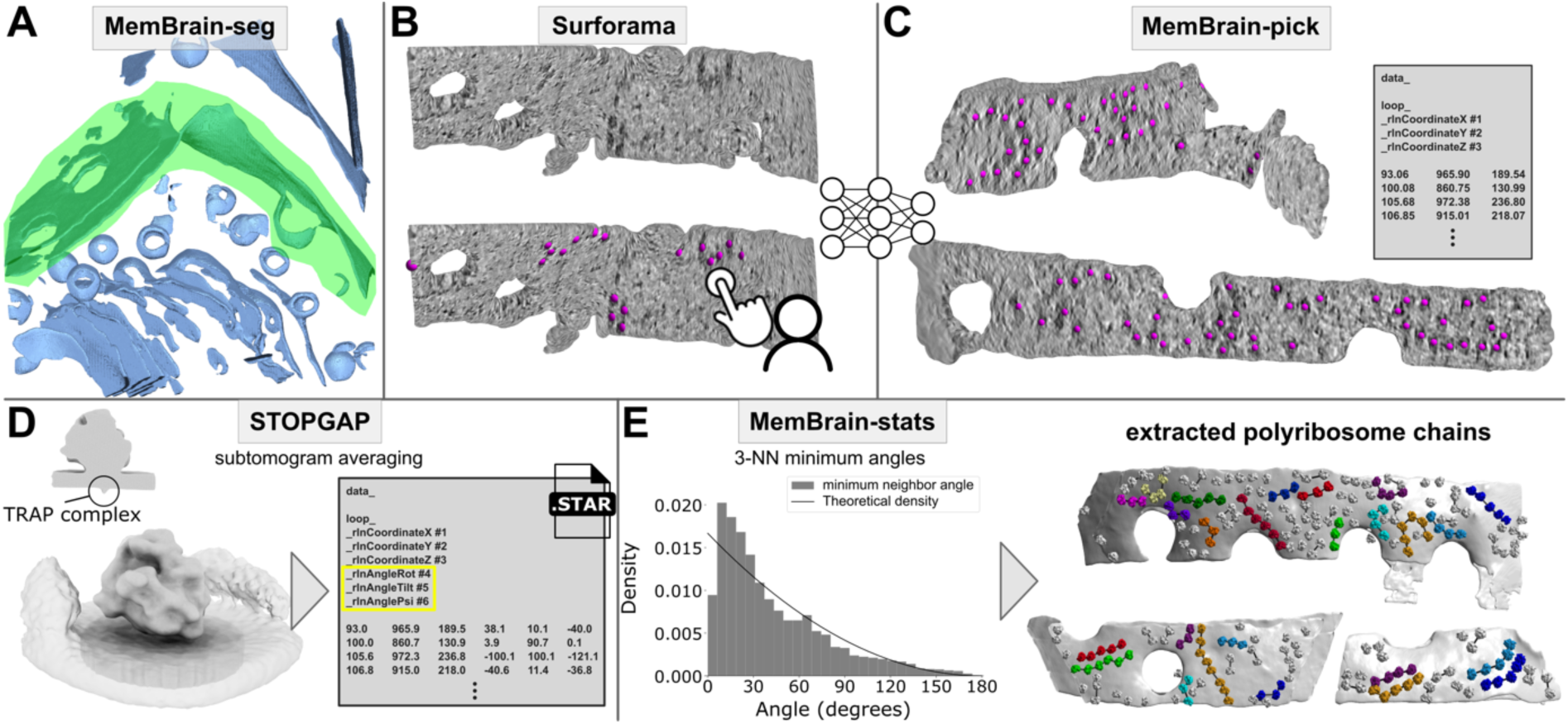
MemBrain v2 end-to-end workflow analyzes nearest neighbor orientations and finds poly-ribosome chains on outer membranes of the nuclear envelope. **A:** MemBrain-seg prediction (light blue) on an entire tomogram from EMPIAR-11830 depicting a nuclear envelope and surrounding membranes. The nuclear envelope is isolated using the MemBrain lasso tool in Napari (green: selected volume). **B:** Top: Visualization of the extracted membrane from A in Surforama. Bottom: GT ribosome positions (magenta) are manually annotated to train MemBrain-pick. **C:** MemBrain-pick predictions (magenta) on another nuclear envelope from the same dataset after training with data from B. **D:** Subtomogram averaging results. Left: subtomogram average depicting a membrane-bound ribosome (22.7 Å resolution from all 4515 particles identified by MemBrain-pick), with a clipped view highlighting the transmembrane TRAP complex. Right: .star file as output giving protein orientations in addition to positions. **E:** Left: Histogram of three-nearest-neighbor (3-NN) minimum angles: For each position, we plotted the minimum angle between the corresponding orientation and the three nearest neighbors’ orientations. The theoretical density depicts the behavior for uniformly random angles between 0 and 180 degrees. Right: Extracted poly-ribosome chains (different colors) using a combination of in-plane angles and distances between nearest neighbors. Grey ribosomes: not part of a chain of length at least 3.

To automate ribosome localization, we manually annotated 13 membranes, trained a MemBrain-pick model, and predicted ribosome positions across the remaining 79 membranes, detecting a total of 4515 positions (example predictions in Fig. 5C). To validate these predictions, we performed subtomogram averaging (STA) in STOPGAP^51^, which resulted in a map clearly depicting a membrane-bound ribosome (Fig. 5D), including density for the transmembrane TRAP complex^74^. The average also contains fuzzy peripheral densities likely corresponding to neighboring ribosomes, which can also be inferred by inspecting the distribution of the picks (Fig. 5E). In addition to structural validation, STA provided protein orientation estimates, enabling further spatial analysis with MemBrain-stats.

To further quantify ribosome organization, we analyzed nearest-neighbor orientation patterns. The distribution of minimum angles among the three geodesic nearest neighbors revealed a shift towards smaller angles compared to a uniformly random orientation model (Fig. 5E left). This result aligns with previous studies reporting ribosome clustering in chains along the nuclear envelope^3,75,76^. Leveraging MemBrain-stats output, we extracted poly-ribosome chains by integrating constraints on nearest-neighbor angles, distances, and chain continuity (Fig. 5E right). This end-to-end MemBrain analysis from raw tomograms to extracted poly-ribosome chains can facilitate the study of ribosome biology in diverse organisms.

## Discussion

The analysis of membranes by cryo-ET is beneficial for many biological studies, as membranes and their associated proteins play central roles in cellular processes. However, the 3D nature of cryo-ET data introduces significant challenges. Manual segmentation of membranes and annotation of their embedded proteins is tedious and time-consuming, particularly because 3D structures require inspection from many orientations and on different planes. Additionally, a fundamental challenge in cryo-ET remains the lack of publicly available GT annotations, limiting the development of robust machine learning models.

MemBrain v2 addresses these challenges by providing an integrated solution for robust membrane segmentation, membrane-associated particle localization, and quantitative spatial analysis. The MemBrain-seg module generates generalizable membrane segmentations using a diverse, well-annotated training dataset and specialized augmentations. By making this dataset publicly available, we provide a foundation for further development and benchmarking of different segmentation methods. To facilitate benchmarking, we include the Surface-Dice metric, which is more sensitive to topological continuity than the traditional Dice score. The MemBrain-pick module offers interactive and data-efficient model training for the localization of membrane-bound particles, significantly reducing the effort required for large-scale analysis. Finally, MemBrain-stats provides quantitative tools for analyzing membrane-bound particle distributions, enabling users to extract meaningful biological insights from native membranes. Together, these components form an end-to-end pipeline for membrane analysis in cryo-ET.

Despite these advancements, certain limitations remain. While MemBrain-seg robustly captures membrane morphology, the resulting segmentation thickness does not necessarily reflect the true biological membrane thickness. Variations in segmentation thickness may stem from annotator-dependent biases, such as differing brush sizes (see Supp. Fig. S7). Another challenge is to maintain MemBrain-pick’s performance when trying to localize diverse membrane protein complexes, as the heterogeneity of targets can complicate universal model performance. Future improvements could involve integrating MemBrain v2 with complementary tools like TomoTwin^56^, to improve versatility across various membrane and protein complex types. For MemBrain-pick, this could lead to more expressive network input features, while MemBrain-seg could also potentially benefit in that this would enable it to distinguish different membrane types in a self-supervised way. Another challenge stems from the inherent constraints of cryo-ET data – feature visibility depends on contrast and noise levels of the tomograms, meaning faint or occluded proteins may be overlooked or poorly annotated. Additionally, strong membrane density signals may dominate and obscure weaker signals from small integral membrane proteins, complicating their accurate detection. These challenges point toward opportunities for future innovation.

MemBrain v2 is built with accessibility and practicality in mind. Its command-line interface for MemBrain-seg simplifies segmentation workflows, making advanced membrane analysis accessible even to users without programming expertise. The seamless integration of MemBrain-pick with Napari plugins fosters an intuitive and interactive experience for protein localization, enabling rapid model refinement and validation. Importantly, MemBrain-seg has already gained significant traction within the cryo-ET community. Many groups have adopted it for their membrane segmentation tasks^12,15,21,22,31,33,77–91^, and it has been applied by the Chan Zuckerberg Imaging Institute to their extensive public cryo-ET dataset^65^. By making all components and datasets open source, MemBrain v2 aims to support collaboration and transparency within the scientific community, offering a versatile and reliable tool for advancing cryo-ET research.

## Data Availability

All tomograms used in this study are publicly available. The latest version of our MemBrain-seg training dataset is accessible via Zenodo: https://zenodo.org/records/15089686

For MemBrain-seg demonstrations, we used tomograms from: CZII-DS-10007, CZII-DS-10224, CZII-DS-10442, CZII-DS-10443, CZII-DS-10444, EMD-3977, EMD-4869, EMD-12329, EMD-12727, EMD-12749, EMD-15407, EMD-16084, EMD-18306, EMD-18748, EMD-30364, EMD-35019, EMD-43050, EMD-44176, EMD-50605, EMPIAR-10988, EMPIAR-11058, EMPIAR-11370, EMPIAR-11830, and EMPIAR-12612.

For MemBrain-pick training and evaluation, we used tomograms from EMPIAR-12612, EMD-10780-10783, EMPIAR-11830, and EMD-10409. To showcase the entire MemBrain v2 workflow, we utilized data from EMPIAR-11830 and EMD-31244.

The GT annotations used for performing the efficiency analysis in MemBrain-pick are available via Zenodo: https://zenodo.org/records/15090084

The Chlamydomonas nuclear envelope-bound ribosome subtomogram average will be deposited under EMD-XXXXX.

## Code Availability

The full MemBrain v2 program is pip-installable via PyPI (pip install membrain) and via https://github.com/CellArchLab/MemBrain-v2.Individual modules modules can be accessed via the following repositories, each of which contains detailed documentation:

- https://github.com/teamtomo/membrain-seg
- https://github.com/CellArchLab/membrain-pick
- https://github.com/CellArchLab/membrain-stats
- https://github.com/CellArchLab/napari-lasso-3d

## Author contributions

Conceptualization by T.P., B.D.E., and L.L, with support from A.M.-S. in the design of Surface-Dice and skeletonization. Development of main functionalities by L.L. Software implementations by L.L., R.D.R., and H.Z., with support from K.Y., A.B, Y.L., S.Zi, and F.I. Data acquisition and initial membrane annotation by W.W. Further patch annotations by S.Zu., L.L., and S.Zi. Testing and suggestion of features by W.W., R.D.R., F.W., and S.Zu. Subtomogram averaging by F.W., and template matching by R.D.R. Supervision by T.P., B.D.E., J.A.S., and A.M.-S. The original draft writing by L.L. Review and editing of the paper by T.P., B.D.E., and R.D.R., with feedback from all authors.

## Acknowledgements

We are grateful to everyone who has tested and provided feedback on the beta version of MemBrain v2, including Matthias Pöge, Robert Brandt, Przemek Dutka, Wangbiao (Seven) Guo, Benoit Gallet, Thomas O’Sullivan, Xiyan Zhu, Zhen Hou, Peijun Zhang, Veijo Salo, Thomas Hoffmann, Giulia Zanetti, Jannis Anstatt, Jenny Sachweh, Abraham (Bram) Koster, Yunjie Chang, Yury Bykov, Pankaj Vilas Jadhav, Duolin Shepherd, and Christos Papantoniou. We thank Abhay Kotecha and his team at Thermo Fisher Scientific for leading the Chlamydomonas visual proteomics project that generated much of the raw data (EMPIAR-11830) used for initial training of MemBrain-seg. Special thanks to users who also contributed training patches and annotations to improve MemBrain-seg’s generalizability: Virly Ananda, Paula Perez Navarro, Benedikt Wimmer, Ohad Medalia, Ya-Ting (Atty) Chang, Michaela Medina, Ben Barad, Danielle Grotjahn, Malit Jessie James Limlingan, Martin Pilhofer, Lena Thärichen, Leanne de Jager, Marten Chaillet, Friedrich Förster, Ben Silva, Donghyun “Raphael” Park, Rory Hennell James, Thomas Marlovits, Sven Klumpe, Jasmina Redzovic, Fridolin Koch, Grigory Tagiltsev, and John Briggs. We are also grateful to Daniil Litvinov for his helpful contributions during MemBrain’s development. These contributions have significantly improved MemBrain-seg’s capabilities. Calculations were performed at sciCORE (https://scicore.unibas.ch/) scientific computing center at the University of Basel. L.L. acknowledges support from the Munich School for Data Science (MUDS) and a fellowship from Boehringer Ingelheim Fonds. A.M-S. is supported by grants RYC2021-032626-I and CNS2023-144921 funded by MICIU/AEI /10.13039/501100011033 and the European Union by NextGenerationEU/PRTR, and the grant PID2023-151075OA-I00 funded by MICIU/AEI/10.13039/501100011033, FEDER and EU. A.M-S. is also supported by grant FSRM/10.13039/100007801(22686/PI/24) funded by Fundación Séneca and the Universidad de Murcia through its program AttractRyC 2023. B.D.E. acknowledges support from an HFSP Research Grant (RGP0005/2021) and ERC consolidator grant “cryOcean” (fulfilled by the Swiss State Secretariat for Education, Research and Innovation, M822.00045). A.B. is supported by the Medical Research Council [MC_UP_1201/6].

## Methods

### MemBrain-seg

MemBrain-seg is a 3D U-Net-based tool designed for generalizable membrane segmentation in cryo-ET data. It incorporates an iteratively generated training set, specialized augmentations, and a Surface-Dice loss function to enhance segmentation accuracy.

#### U-Net

##### Architecture

MemBrain-seg’s U-Net architecture is based on design choices from nnU-Net^92^, a leading biomedical image segmentation framework, and is implemented using MONAI^93^. It consists of five downsampling and upsampling blocks, with deep supervision^94^ to improve gradient flow and convergence.

##### Loss function

We use a composite loss function combining binary cross-entropy, Dice loss, and Surface-Dice loss (see Section <u>“Surface-Dice”</u>). To handle uncertain image regions, an ignore feature excludes certain voxels from loss calculations (see Section <u>“Iterative Dataset Generation”</u> and Supp. Fig. S3).

##### Training setup

All models were trained for 1000 epochs using stochastic gradient descent, a batch size of 2, and 160^3^-shaped training patches on RTX 4090 GPUs. We apply a polynomial learning rate scheduler decreasing from 0.01 to 0.0. Before training, patch intensities are normalized per-patch by subtracting the mean and dividing by the standard deviation.

##### Inference

For inference, tomograms are processed using a sliding window approach with Gaussian-weighted patch aggregation of scores. Each tomogram is divided into overlapping 160^3^ patches, and we apply 8-fold test-time augmentation by flipping along all axis combinations, averaging predictions before thresholding. MemBrain-seg also supports internal rescaling to a specified pixel size (default 10Å). In this mode, patches are extracted at the original resolution, rescaled before passing through the network, then restored to the original scale before aggregation. As rescaling occurs entirely on GPU, this approach significantly speeds up processing compared to full tomogram rescaling.

#### Data augmentations

Data augmentation is critical for improving generalization by simulating the diverse appearances of tomograms and membranes^95^. Below, we describe the geometric, intensity-based, and Fourier-based augmentations applied during training.

##### Conventional data augmentations

We apply a mix of classically used geometric and intensity transformations. These augmentations, applied randomly and on-the-fly, introduce strong variation in training samples, enhancing the network’s adaptability (see Supp. Fig. S8A).

*Geometric Transforms:* These transformations aim to show the image from different viewing points, and include rotation at arbitrary angles, zooming within a range of 0.7 to 1.4, and both shuffling and flipping of axes.

*Intensity Transforms.* Applied to alter image characteristics, these encompass median filtering, Gaussian blurring, Gaussian noise addition, brightness and contrast adjustments, low-resolution simulation, random erasing, additive brightness gradient, local Gamma transform, and sharpening.

##### Fourier Amplitude Augmentation

Fourier Amplitude Spectrum Matching, as introduced in DeePiCt^44^, adjusts tomogram styles by matching their frequency-domain amplitudes. It does so by rotationally averaging Fourier intensities per frequency band, generating a 1D sequence for both input and target tomograms. The ratio of these sequences is used to create a 3D radial filter, which is then multiplied in Fourier space to transform the input tomogram.

In MemBrain-seg, however, our goal is to improve generalization without requiring explicit style normalization before inference. Previous studies have shown that training with style augmentations is more effective for generalization than pre-prediction normalization^96,97^ . We therefore introduce Fourier Amplitude augmentation, which randomly modifies tomogram styles during training. Rather than deriving transformations from a limited set of reference tomograms, we generate random normalization sequences to simulate diverse styles dynamically.

To achieve this, we create a 1D sequence x of length 80 (i.e., half the patch size) using a random walk:

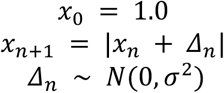

where σ = 0.2, | . | is the absolute value, and *N* is a normal distribution. This sequence is expanded into a 3D radial kernel and multiplied with the Fourier transform of the input volume, introducing random style variations. The effects on image appearance are illustrated in Supp. Fig. S8B: Up-scaling of higher frequencies results in noisier images with weaker membrane contrast (2nd row), whereas down-scaling of high frequencies enhances coarse structures, making membranes more prominent (3rd row).

##### Missing Wedge augmentation

Segmenting membranes in regions affected by the missing wedge presents a significant challenge due to the anisotropic distortions it introduces. To mitigate this, we adopt an approach inspired by IsoNet^40^, in which we rotate input subvolumes randomly before applying an artificial missing wedge by masking Fourier coefficients in a wedge-like shape. This controlled simulation (Supp. Fig. S8C) replicates the characteristic loss of densities caused by incomplete angular sampling in cryo-ET. In experimental data, the information lost due to the missing wedge is not directly visible, yet in some cases, the image context provides hints about where membranes should be. By artificially replicating these effects, we train the network to infer missing structures based on contextual information. Training pairs are generated by applying this transformation to subvolumes while preserving their unaltered ground truth (GT) labels, pushing the model to learn to accurately segment membranes even in wedge-impacted regions.

#### Surface-Dice

Surface-Dice extends Centerline-Dice^98^ to 3D membrane segmentation, serving as both a metric for evaluating binarized membrane segmentation outputs and a loss function during network training. By leveraging a differentiable skeletonization approach, Surface-Dice better reflects the continuity and consistency of membrane structures in 3D than normal Dice scores.

##### Computation

Similar to Centerline-Dice, Surface-Dice relies on skeletonizations of both the predicted segmentation (Mpred) and the ground truth (MGT), denoted as Spred and SGT, respectively. Skeletonization in Surface-Dice reduces the membrane segmentation to a 1-voxel-thick surface, forming a 2D manifold in 3D space (illustrated in Supp. Fig. S6A,B). To compute Surface-Dice (Dicesurf), we define Surface-precision (Precsurf) and Surface-recall (Recsurf) as follows:

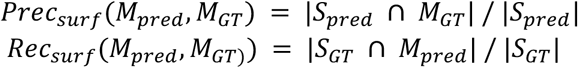

The |·|-operator sums up all positive voxels in the respective set. Surface-Dice (Dicesurf) is then computed as the harmonic mean of these two terms:

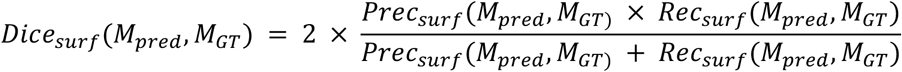

##### Surface-Dice as a Loss Function

To use Surface-Dice as a loss function during training, we avoid thresholding network outputs, as this operation is non-differentiable. Instead, we employ a differentiable skeletonization approach, similar to the method used in Centerline-Dice. This process involves iterative membrane erosion to progressively thin the segmentation. At each iteration, an additional erosion and dilation step is applied, and differences between the segmentation before and after these operations are evaluated. The presence of differences indicates that erosion has removed portions of the membrane, meaning the current iteration already represents a thin surface in those regions. Both erosion and dilation are implemented via min- and max-pooling, ensuring that all operations remain differentiable and efficient. For further details, please refer to the Centerline-Dice publication^98^.

#### Iterative Dataset Generation

To efficiently build a well-annotated membrane segmentation dataset, we employed an iterative approach inspired by active learning principles^99^. This strategy minimized manual annotation efforts by focusing on areas where the network struggled with segmentation. The schematic correction workflow and different iterations are visualized in Fig. 2B and <u>Supp. Fig.</u> <u>S3</u>. During the annotation process, we pay particular attention to the quality of the training dataset, i.e., the accuracy of the segmentation. The ignore label is particularly helpful here, as it prevents the model from learning uncertain regions simply as non-membrane background. This annotation quality is a key factor for the generalization capability of our network. Detailed instructions on how users can annotate training patches to customize the model can be found in our online documentation: https://github.com/teamtomo/membrain-seg/blob/main/docs/Usage/Annotations.md

##### Initial Dataset and First Iteration

We initiated our project with membrane segmentation patches from the *Spinacia oleracea* dataset, initially annotated using TomoSegMemTV^42^. Using MITK Workbench^100^, we manually refined these patches, each sized 160^3^ voxels, assigning to every voxel either the background, membrane, or ignore classes – the latter not being evaluated by the loss function during training as it marks uncertain membrane regions, where exact delineation of membranes was challenging. The process of correcting predicted membrane segmentations is depicted in Supp. Fig. S3B, where we remove false positive voxels from the segmentation, add falsely negative voxels to it, and assign the ignore class in uncertain regions. Using these corrected annotations, we trained an initial U-Net^43,92^ model for membrane segmentation. While this model outperformed TomoSegMemTV, its segmentations remained suboptimal. We therefore conducted a second round of annotations, targeting patches where the model’s predictions were weak. This refinement process resulted in 69 accurately annotated spinach chloroplast patches (<u>Supp.</u> Table 1, column “Spinach”; Supp. Fig. S3B, “Round 1”).

##### Expansion to Further Datasets

To improve generalization across different imaging setups, we extended our approach to *Chlamydomonas reinhardtii* tomograms from EMPIAR-11830^73^. Due to differences in imaging conditions, initial segmentations performed worse than in the spinach dataset. By iteratively re-annotating 33 patches (covering thylakoid membranes, mitochondria, and the Golgi apparatus), we aimed to improve performance on this dataset (<u>Supp.</u> Table 1, column “Chlamy”; Supp. Fig. S3B, “Round 2”). To further expand the model’s applicability beyond our own data, we collaborated with other research groups who tested MemBrain-seg on their own datasets and provided re-annotated patches from regions where MemBrain-seg struggled. After careful quality assessment, these contributed 27 additional patches, significantly increasing dataset diversity (<u>Supp.</u> Table 1, column “Community”; Supp. Fig. S3B, “Round 3”). This ongoing collaboration continues to improve MemBrain-seg’s performance across a wide range of tomograms. Instructions on how to annotate and share patches can be found on the MemBrain-seg Github repository.

##### Adding Publicly Available Sources

We further leveraged the few existing publicly available segmentation sources to improve our model even further. To achieve this, we generated 40 membrane patches using the open-source tomography simulators PolNet^66^ and CryoTomoSim^39^ (20 patches each; <u>Supp.</u> Table 1, column “Synthetic”; Supp. Fig. S3B, “Round 4”). Additionally, we integrated an externally annotated dataset by extracting 15 reliable segmentation patches from the publicly available DeePiCt dataset^44^ (EMPIAR-10988). These regions were selected from areas where MemBrain-seg initially struggled, enhancing dataset diversity (<u>Supp.</u> Table 1, column “DeePiCt”; Supp. Fig. S3B, “Round 5”).

#### Skeletonization

MemBrain-seg provides a functionality to convert binary segmentations into 1-voxel thick membrane sheets, which is essential for several downstream applications and the computation of Surface-Dice. We solve this task similarly to TomoSegMemTV^42^: First, we compute a distance transform (DT), converting the segmentation into a distance map relative to membrane boundaries. The goal is to identify the center sheet, which corresponds to the regions with the highest distance values. To extract this center sheet, we approximate membrane normal vectors by calculating the Hessian matrix of the distance transform and extracting its eigenvectors at each voxel p. The eigenvector n associated with the largest intensity curvature provides the direction pointing outward from the membrane. We then apply non-maximum suppression (NMS) along these normal vectors: A voxel is retained if it has the highest DT value along its normal vector; otherwise, it is suppressed:

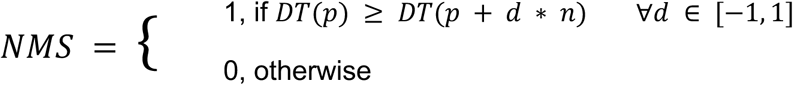

This process yields a 1-voxel-thick sheet, representing the center sheet of the membrane segmentation.

#### Model Fine-tuning

MemBrain-seg’s segmentation performance can be affected by batch shifts, leading to suboptimal results when tomogram or membrane appearances deviate significantly from the training domain. To mitigate this, we implemented a fine-tuning strategy that allows users to quickly adapt the model to their specific datasets. Fine-tuning requires a small number of annotated patches (typically 10–20, each 160^3^ voxels at 10Å pixel size) and follows a transfer learning approach^101^, where pretrained model weights serve as initialization. To prevent overfitting and reduce computational costs, we limit training to 100 epochs (instead of 1000) and lower the learning rate to 10^−5^ from 10^−2^. An early stopping criterion halts training if validation performance deviates too far from the original model’s performance, ensuring stability. To enable this, the original, full validation dataset serves also as validation dataset in this task.

For fine-tuning, incorporating the Surface-Dice loss can be beneficial, as it disregards variations in annotation thickness (e.g., Supp. Fig. S7), preventing premature stopping due to segmentation thickness differences. This strategy enables MemBrain-seg to efficiently adapt to new datasets, enhancing performance across diverse tomographic conditions. The workflow is illustrated in Supp. Fig. S9.

#### MemBrain-seg usability

MemBrain-seg is designed to be as easy to use as possible. The default way of using it is by a command line interface. Via its API, membrain-seg is also easy to integrate into other software packages and existing pipelines, allowing subsequent analysis of membrane segmentations. Additionally, MemBrain-seg is already integrated into other packages with a graphical user interface, like ColabSeg^102^ and Scipion^103^.

### MemBrain-pick

MemBrain-pick enables the automated localization of membrane-associated proteins in cryo-electron tomograms. Unlike other deep learning approaches, it operates directly on membrane surfaces rather than voxel-based representations, allowing for a more specialized and focused detection of membrane-bound proteins.

#### Workflow

The MemBrain-pick workflow consists of three key steps: data preparation, model training, and prediction. Data preparation consists of extracting relevant membrane areas from MemBrain-seg segmentations, and conversion to mesh representations. The model is then trained with membrane meshes as input, where we project tomographic densities onto the mesh triangles (see Fig. 3, Supp. Fig. S10). We train the DiffusionNet-model^67^ to predict a membrane particle distance map. During inference, we extract particle positions using Mean Shift clustering^104^.

##### Data Preparation

#### Mesh generation and projection

To represent membranes as analyzable surfaces, we first extract membrane segmentations from MemBrain-seg (e.g. crop using our Lasso tool, see Section <u>“Napari</u> <u>Tools”</u>) and convert them into triangular mesh representations. This is achieved using the Marching Cubes algorithm^105^. To ensure a uniform mesh resolution, we resample triangle sizes through Voronoi clustering^106^, which redistributes vertices to achieve evenly sized barycentric areas per vertex. Once the mesh is generated, we leverage its per-vertex normal vectors to extract relevant tomographic information. For each vertex, we sample intensity values along its normal vector in both directions, generating a feature vector of N sampled density values per vertex.

#### Training data generation

MemBrain-pick requires GT annotations for training. To facilitate accurate annotation, MemBrain-pick is compatible with Surforama^60^, a Napari-based^64^ annotation tool that operates similarly to MemBrain-pick by displaying tomographic densities projected onto a membrane surface. Using Surforama, users can efficiently annotate protein center positions directly on membrane meshes. These annotations are exported as STAR files^107^, ensuring seamless integration with MemBrain-pick’s training pipeline, as well as other processing software.

#### Surface partitioning

To optimize model efficiency and prevent overfitting to global membrane geometries, we partition each membrane into smaller overlapping surface crops before feeding them into MemBrain-pick. Each partition consists of 2,000 triangles, selected by initializing at a random seed triangle and iteratively expanding to include the 2,000 nearest neighbors based on geodesic distance. To ensure comprehensive coverage, partitions are generated with sufficient overlap so that each mesh vertex has a neighboring seed within close proximity. This procedure maintains enough in-plane context per-patch for accurate protein localization while reducing computational complexity and avoiding learning global geometries.

##### Training

#### Training objective from GT

Rather than directly predicting protein center locations, MemBrain-pick is trained to estimate a continuous distance field relative to the nearest GT protein position. For each mesh vertex, we compute the Euclidean distance to its closest annotated protein center, generating a smooth distance map that serves as the training target. The network is optimized using a Mean Squared Error (MSE) loss, minimizing the discrepancy between predicted and GT distance values.

#### Data Augmentations

To enhance generalization and prevent overfitting, we developed a suite of point cloud augmentations to be applied during training. These augmentations introduce controlled variations in spatial and feature domains, mimicking fluctuations in tomographic appearance:

- *Spatial Gaussian Smoothing*: Smooths point cloud features by computing a weighted average of neighboring points, with Gaussian weights based on spatial distance and a randomly drawn sigma.
- *Feature Gaussian Smoothing*: Applies a Gaussian filter along the normal vector direction to reduce noise in per-vertex features.
- *Feature Dropout*: Randomly sets a fraction of features to zero.
- *Feature Noise*: Adds Gaussian noise to each feature to simulate signal variability in tomographic data.
- *Feature Shift*: Offsets feature values by adding a random constant, introducing brightness shifts.
- *Feature Scale*: Scales feature intensities by a random factor to account for contrast variations.
- *Random Erasing:* Selects small patches in the sample and zeroes out all features within them.
- *Random Brightness Gradient*: Scales feature intensities proportionally to the dis tance from a randomly chosen point within or near the sample.
- *Random Brightness Gamma*: Applies a gamma correction, where the gamma value is determined based on distance from a randomly sampled point.

Effects of combinations of these augmentations during training are visualized in <u>Supp. Fig.</u> <u>S16</u>.

#### Network Architecture

We use DiffusionNet^67^, a deep learning framework designed for learning on mesh surfaces. DiffusionNet propagates per-vertex features using a learned diffusion process, incorporating both geometric and contextual information. We apply it to membrane meshes, where input features are derived from tomographic intensity projections, and training targets are distance maps representing protein localization. To enhance performance, we integrate a learnable 1D convolutional layer that processes features along the membrane normal before diffusion. This is inspired by the concept of separable convolution^108^. Additionally, we stack multiple DiffusionNet blocks to capture both local curvature and broader membrane context.

##### Inference

#### Merging Overlapping Partitions

Each partition is processed independently, producing per-vertex protein distance maps. To ensure seamless predictions, we assign each vertex a Gaussian-decaying weight from the partition seed and merge overlapping predictions using a weighted average, similar to a sliding window approach. This reduces edge artifacts and ensures continuity across partitions.

#### Membrane Normal Distance Shifts

To compensate for segmentation inaccuracies, we perform inference multiple times with shifted feature extraction windows. Instead of using a single fixed range along the membrane normal, we iteratively shift the sampled density range in small steps. The final prediction is obtained by averaging across shifts, improving robustness to variations in membrane thickness and segmentation offsets.

#### Mean Shift Clustering

After generating protein center distance maps with DiffusionNet, we apply Mean Shift clustering^104,109^ to extract protein center locations. First, we threshold predicted distances to remove clear background regions. For the remaining vertices, we refine cluster centers using a score-guided adaptation of Mean Shift, similar to [63], where each vertex p updates its position based on a weighted average of its neighbors:

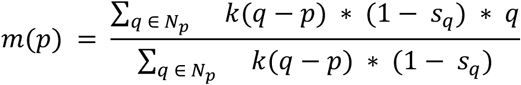

Here, Np includes all points within a radius b around p, excluding p itself, and sq represents the normalized predicted distance score. The bandwidth b, defining the neighborhood size, is set to match the maximum allowed distance score in our training data. We use Euclidean distance as the kernel function k, and the additional weighting by predicted scores to improve convergence towards high-confidence protein centers.

To accelerate clustering, we use a PyTorch-based GPU implementation^104^, significantly reducing inference time. Once mean shift has converged, we apply density-based spatial clustering (DBSCAN^110^) to merge redundant cluster centers and eliminate outliers, refining the final protein localizations.

### MemBrain-stats

#### Particle Concentration

Particle concentration is computed as the number of detected particles divided by the total surface area of the membrane mesh. For analyses focused on a single membrane side, we provide two approaches to ensure accurate estimation: either by dividing the total area by two, assuming particles appear only on one side and both sides are of roughly equal size (i.e. not much membrane curvature), or by isolating a single membrane side surface through edge exclusion (see Section <u>“Edge Exclusion for Robust Analysis”</u>), followed by connected component filtering.

#### Geodesic Distance Calculation

Geodesic nearest neighbors are computed using either an exact shortest-path algorithm for triangular meshes^111^ or an approximate method based on the heat diffusion equation^112^. The latter provides a computationally efficient alternative by leveraging the heat method to estimate geodesic distances. In practice, both approaches yield nearly identical results, with the heat method offering a significant speed advantage.

#### Geodesic Nearest Neighbors

To compute geodesic nearest neighbor distances, each detected protein center is first projected onto its nearest mesh vertex. Accurate distance estimation requires a sufficiently fine-grained mesh to minimize quantization errors. Once projected, the N nearest geodesic neighbors are determined for each protein center using one of the above methods.

#### Ripley’s statistics

To compute Ripley’s statistics, we follow the framework described in PyOrg^63^, adapting the formulation to 2D geodesic distances on membrane surfaces, which are modeled as 2D manifolds in 3D space. For a set of membrane particle locations S and a set of triangles T forming the membrane mesh, we define:

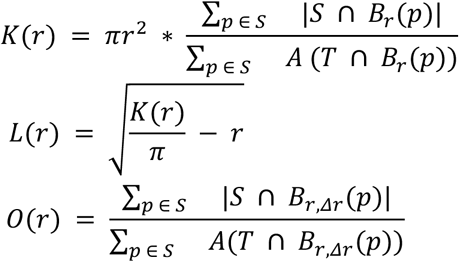

where |.| denotes the number of particle in a given region, A(.) computes the area of triangles in a set, Br(p) is the geodesic ball of radius r around particle p, and B_r_,_Δr_(p) geodesic ring around p, extending from radius r to r + Δ*r*:

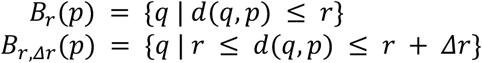

where d(.,.) represents the geodesic distance between two points on the membrane surface.

By restricting area computations to the triangular mesh T, we automatically normalize protein concentration based on the connected membrane surface rather than the entire tomographic volume. As in PyOrg, the functions K, L, and O are closely related: L stabilizes the variance of K, and O can be understood as the derivative of K, providing information on local protein clustering patterns.

#### Edge Exclusion for Robust Analysis

Particle localization can become unreliable near the boundaries of membrane segmentations due to the lack of in-plane context. Additionally, segmented membranes are often truncated at arbitrary heights, meaning that segmentation edges do not necessarily correspond to actual membrane boundaries. This can distort spatial metrics such as nearest neighbor distances. To mitigate these issues, we aim to automatically exclude edge regions from certain analyses.

The marching cubes algorithm gives us triangular meshes depicting the segmentations’ hulls. Since membranes are thin sheets, their edges are represented by regions with high curvature. We detect these areas by computing the discrete mean curvature at each vertex, defined as the average angle between all neighboring triangles within a geodesic sphere of radius R. Membrane edge regions are then approximated by selecting the top 5% of vertices with the highest curvature values. Finally, all points within a distance *d* from these edge vertices are excluded from the analysis to prevent artifacts in localization and spatial measurements. An example of the edge exclusion is visualized in Supp. Fig. S14I.

#### In-plane Orientation Comparison

To analyze the relative in-plane orientations of nearest-neighbor particles, we first compute geodesic nearest-neighbor pairs as described above. We assume that particle orientations are known, for instance, from subtomogram averaging, where each particle is aligned such that its z-axis is parallel to the local membrane normal.

To express the particle’s orientation in the membrane plane, we construct a rotation matrix that aligns the standard unit vector (0, 0, 1)^T^ with the membrane normal at the particle’s location. This rotation matrix transforms the basis vectors, mapping (0, 1, 0)^T^ into the membrane plane, providing a reference direction for in-plane orientation.

For a pair of particles, we compare their in-plane orientations by computing the angle θ between their respective in-plane vectors:

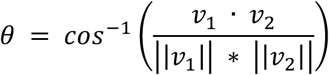

where v1 and v2 are the in-plane orientation vectors of the two particles, · represents the dot product, and]|. |] is the vector’s L2-norm. This provides a quantitative measure of orientation similarity within the membrane plane.

### Napari tools

#### Compatibility with Surforama

MemBrain-pick is fully compatible with Surforama, a Napari-based^64^ tool for annotating membrane proteins in cryo-ET. Similar to MemBrain-pick’s network input, Surforama projects tomographic densities onto the membrane mesh, allowing users to navigate along the membrane normal vector to visually inspect the in-plane densities of embedded particles. Surforama exports annotations in STAR format^107^, which also serves as the preferred input format for training MemBrain-pick. To ensure seamless integration, MemBrain-pick outputs are made compatible with Surforama by generating both STAR files and .h5 containers, which can be directly visualized in Surforama with a single MemBrain-pick command.

#### Lasso functionality (Lasso, masking, connected components)

We developed a MemBrain-pick Napari^64^ plugin to facilitate the transition from MemBrain-seg outputs (full tomogram segmentations) to MemBrain-pick inputs (segmented single-membrane instances). The plugin includes a lasso functionality, allowing users to interactively select and extract 3D regions of interest. Following selection, a connected component analysis is performed to isolate individual membrane segments, which can then be saved for further processing. Additionally, users can refine segmentations using standard Napari annotation tools, enabling cleaner and more precise membrane instance selection. For example use cases, see Supp. Fig. S1B-E.

### Applications

#### MemBrain-seg experiments

All membrane segmentations shown in Fig. 2, Supp. Fig. S2 were generated using MemBrain-seg’s `membrain segment` command using the `MemBrain_seg_v10_alpha` model. Visualizations were generated using ChimeraX^113^.

#### MemBrain-seg Ablation Studies

To assess the impact of different dataset components, we conducted a five-fold cross validation experiment (see <u>Supp.</u> Table 1). Each fold consists of patches from distinct tomograms, ensuring no tomogram appears in both the training and test sets at the same time. Given the limited size of our dataset, this cross-validation strategy provides a more robust evaluation of model performance.

For each iteration, the model was trained using four folds and evaluated on the remaining one, cycling through all folds so that each served as the test set once. Performance was measured using the Surface-Dice and standard Dice score, offering a comprehensive assessment of segmentation quality. In the plots (Supp. Fig. S5), we report the mean and standard deviation across all folds, along with individual run results.

##### Incremental Training

To evaluate how increasing dataset diversity enhances MemBrain-seg’s performance and generalization, we incrementally trained models by sequentially incorporating the five rounds of annotations as detailed in Section <u>“Iterative Dataset Generation”</u>. Each step followed the same cross-validation setup described above. The results of this incremental training approach are presented in Supp. Fig. S5A.

##### Data Augmentations

To analyze the effect of Fourier-based data augmentations, we trained models using the dataset from incremental training round 4, i.e., excluding the DeePiCt data. By training on this subset, we specifically examined the impact of augmentations on generalization to the DeePiCt test set, serving as an external generalization dataset. The results, are shown in Supp. Fig. S5B, highlighting the improved performance with augmentations.

##### Fine-tuning with Surface-Dice

To evaluate the performance of fine-tuning, we initialized models using those trained in incremental training round 4 and fine-tuned them with round 5 data (DeePiCt) while ensuring fold separation remained consistent. We tested two fine-tuning strategies: one using Dice and BCE loss for optimization and validation tracking, and another incorporating Surface-Dice into the loss function. The impact of these strategies is shown in Supp. Fig. S9B,C.

#### TARDIS Comparison

To compare MemBrain-seg with TARDIS, we downloaded the latest available TARDIS model for 3D membrane segmentation (download date: 6th February 2024). We applied the model to all our patches using the `tardis_mem` command to obtain binary segmentations for each patch. For the evaluation presented in Supp. Fig. S5A, we computed Dice and Surface-Dice scores relative to our GT, excluding regions labeled as “ignore”.

It is important to note that our GT dataset was generated iteratively, incorporating incremental versions of MemBrain-seg. Although we manually corrected segmentation errors, the dataset still exhibits some bias toward MemBrain-seg predictions. As such, direct comparisons with other programs should be interpreted with caution.

#### MemBrain-pick quantitative evaluation

##### Evaluation Metric

To make results comparable with metrics reported in MemBrain v1^61^, we also adopted the same calculation of F1-Scores to quantify the MemBrain-pick performance: Given a radius r, for predicted position, we define it a true positive (TP_p_) if it is within distance r of a GT position, and a false positive (FPP) otherwise. Similarly, for a GT position, a (TPR) is within distance r or a predicted position, and a (FNR) is not. With this, we can define recall, precision, and the F1-Score (harmonic mean of precision and recall):

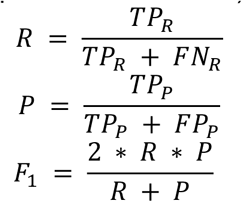

In our experiments, we chose the same radius also used in [63], which is 4.5 bin-4 voxels, corresponding to 4.5 * 14.08Å = 63.36Å.

##### Spinach data

The Spinach dataset used to evaluate MemBrain-pick’s performance is described in more detail in a separate publication^19^. To train the MemBrain-pick model used in Fig. 3C, we used the same dataset as in MemBrain v1^61^, consisting of 45 membrane segmentations from nine tomograms of *Spinacia oleracea* thylakoid membranes. Previously existing GT annotations used in MemBrain v1 were manually refined to improve accuracy and to classify particles into three categories: Photosystem II (PSII), Cytochrome b6f (b6f), and unidentifiable densities (UK).

Following the protocol in MemBrain v1, we designated all ten membranes from two tomograms as the test dataset. For the incremental training performance plot in Fig. 3D, five membranes from the remaining 35 (fixed across runs) were assigned to the validation set. Training sets were constructed by randomly sampling N membranes from the remaining 30, where for each value of N, five independent subsets were drawn, and a separate model was trained for each subset. The same dataset splits were used for training both MemBrain-pick and MemBrain v1 models.

#### Chlamy Generalization

To evaluate the generalization performance of MemBrain-pick, we applied a model trained exclusively on *Spinacia oleracea* data to two independent *Chlamydomonas reinhardtii* datasets, acquired under different imaging conditions but both depicting thylakoid membranes with embedded PSII and b6f complexes (Fig. 3E, F). Details on the acquisition and setup of the Chlamyold^17^ and Chlamynew^73^ datasets can be found in their respective publications.

For Chlamyold, we used 28 segmented membranes extracted from four tomograms, along with GT annotations from the original publication^17^, enabling the computation of F1-Scores (Fig. ^3E^ inset). The Spinach-trained MemBrain-pick model was applied directly to these membranes to predict particle positions.

For Chlamynew, we first segmented all membranes in four tomograms by applying MemBrain-seg to Cryo-CARE-denoised^36^ tomograms. Using the MemBrain Napari plugin, we extracted 23 thylakoid membranes from these full-tomogram segmentations. To further enhance image interpretability, we applied IsoNet^40^ before predicting particle positions with the Spinach-trained MemBrain-pick model (Fig. 3F).

#### Phycobilisomes

For the analysis presented in Fig. 4, we used data from red algae^71^, which depicts phycobilisome-photosystem II supercomplexes (EMD-31244). Membrane segmentations were generated using MemBrain-seg (`MemBrain_seg_v10_alpha` model), followed by connected component splitting via the MemBrain Napari plugin, yielding 29 individual membrane segmentations. To provide GT annotations, we manually labeled phycobilisome positions on five membranes using Surforama and trained MemBrain-pick with its default parameters. The trained model was then applied to the remaining 24 membranes in the tomogram, enabling us to compute Ripley’s statistics on this dataset. For computing Ripley’s O statistic, we divided all distances into bins of size 5 nm.

#### Ribosome Analysis in EMPIAR-11830

##### MemBrain-pick prediction

For the ribosome analysis shown in Fig. 5, we extracted particle positions from the *Chlamydomonas reinhardtii* dataset (EMPIAR-11830^73^), focusing on the outer membrane of the nuclear envelope and the endoplasmic reticulum. To scale up the analysis, MemBrain-seg was applied to 140 tomograms (MemBrain_seg_v10_alpha model), extracting the largest connected component from each segmentation. After removing segmentations containing membrane structures of other classes or excessive noise, 97 tomograms remained for further processing. Ribosome GT annotations were manually assigned to 13 membranes using Surforama, and MemBrain-pick was trained with its default parameters. The trained model was then used to predict ribosome positions across the remaining membranes, yielding 4515 predicted positions. Both MemBrain-seg and MemBrain-pick were applied to Cryo-CARE-denoised^36^ tomograms to enhance contrast and improve localization accuracy.

##### Subtomogram Averaging

The endoplasmic reticulum- and nuclear envelope-bound ribosomes from the MemBrain-pick predictions were further processed in subtomogram averaging. Averaging was performed in STOPGAP^51^. Bin4 3D CTF-corrected tomograms were generated as implemented in IMOD^114^, from which subtomograms for all ribosome positions were extracted with a box size of 80 pixels. Since all particles were already pre-aligned normal to the membrane from MemBrain-pick, initial alignment was performed using a so-called “cone-search” which corresponds to a global search around one axis (perpendicular to the membrane) while restricting the search for the other 2 angles. After initial alignment, all angular searches were iteratively reduced as resolution increased, reaching 22.7 Å resolution after 14 iterations. The Fourier Shell Correlation plot using a soft spherical mask is visualized in Supp. Fig. S17.

##### Ribosome Chains

For the analysis shown in Fig. 5E, we first computed the three nearest neighbors for each ribosome and determined the in-plane angles between them using MemBrain-stats, based on orientation estimates from subtomogram averaging. We plotted the smallest of these three angles for each ribosome position and compared it to a theoretical distribution. The theoretical distribution was generated by randomly sampling 100,000 triplets of uniformly distributed angles between 0 and 180 degrees and plotting their distribution. Next, we extracted ribosome chains by iteratively adding chain units based on three spatial constraints: (1) a center-to-center distance below 40 nm, (2) an in-plane angle below 60 degrees, and (3) an angle between the current chain end-connection and a potential new end-connection below 80 degrees. Using these criteria, we identified and reconstructed the ribosome chains shown in Fig. 5E. An example jupyter notebook with showing the extraction of these ribosome chains can be accessed on the MemBrain-stats Github repository.

Note: we used three nearest neighbors instead of only the first nearest neighbor, as the closest ribosomes often did not appear to be part of the same chain (see Fig. 5E, right). This approach allowed for a more reliable identification of polysome-like arrangements.

#### Ribosomes in EMD-10409

To generate the visualization of ER-bound ribosomes in Supp. Fig. S11B, we processed the tomogram from EMD-10409^4^ using MemBrain-seg and MemBrain-pick. All membranes in the tomogram were first segmented using MemBrain-seg (`MemBrain_seg_v10_alpha` model). The segmented membrane was then divided into six non-overlapping regions using the MemBrain Napari tool. Ribosome GT annotations were manually assigned to three of these regions in Surforama, which were subsequently used for training MemBrain-pick. The trained model was then applied to the remaining three regions to predict ribosome positions, as shown in Supp. Fig. S11B. For MemBrain-seg, the raw tomogram was used directly, whereas for MemBrain-pick, we applied IsoNet^40^ to enhance contrast and improve protein localization.

### Respirasomes

To evaluate MemBrain-pick’s performance on mitochondrial membrane-associated complexes, we analyzed respirasomes in *Chlamydomonas reinhardtii* mitochondria in tomograms from EMPIAR-11830^73^. Membrane segmentations were generated from four tomograms using MemBrain-seg (`MemBrain_seg_v10_alpha` model), and 45 individual cristae segmentations were extracted using the MemBrain Napari plugin. To provide GT particle positions, we used previously generated template-matching-based coordinates^20^, projected them onto the membrane surfaces, and used these as training data for MemBrain-pick. The model was trained on 35 membranes from three tomograms and then applied to the remaining ten membranes from the fourth tomogram (shown in Supp. Fig. S11A). All tomograms were denoised using Cryo-CARE^36^ prior to membrane segmentation and particle localization.

### MemBrain-pick comparisons to other methods

In Fig. 3 and Supp. Fig. S13, we compare the performance of MemBrain-pick with other particle picking methods commonly used in cryo-ET. Below, we provide details on the applications of these methods using our Spinach dataset (see Section <u>“Spinach data”</u>).

#### Template Matching (PyTOM)

Template matching (TM) was performed using `pytom-match-pick` ^50^ on the two test set Spinach tomograms. The single-particle cryo-EM map of spinach PSII (EMDB-6617) ^115^ was chosen as a template. The template was resampled to a pixel size of 14.08 Å to match the tomogram’s, and cropped to a box size of 32 pixels. Finally, the contrast was inverted to match that of the tomograms (protein density being black). A slab mask for the tomogram was created using Slabify^116^. The program `pytom_match_template.py` was run using a fan (per-tilt) wedge model, a particle diameter of 310 Å (corresponding to the longest axis of PSII), a high-pass filter of 400 Å, C2 symmetry and random phase correction. The angular sampling was calculated automatically from the particle diameter and pixel size, corresponding to 4.89 degrees along Z1 and X and 5.14 degrees along Z2 (Euler angles in PyTOM conventions).

For extraction, the score maps were masked with individual single membrane masks of the test membranes. Automated extraction using `pytom_extract_candidates.py` with the `--number-of-false-positives` option failed to provide meaningful results, so a custom procedure was adopted. For each membrane, the threshold was calculated by finding the 99th percentile of the scores within the mask. This roughly corresponds to the noise threshold when sorting all the scores within the mask (see Supp. Fig. S13D). A script for this calculation (`tm_find_thresh`) is provided at https://github.com/CellArchLab/cryoet-scripts. Then `pytom_extract_candidates.py` was used to extract up to 100 candidates above this threshold per membrane, with an exclusion radius of 6 pixels (84.48 Å) around each peak, corresponding to the shortest axis of PSII.

#### DeepFinder

To train DeepFinder^54^, we performed a training-validation split at the tomogram level to prevent overlapping patches within a single tomogram. From the total nine tomograms, five tomograms were used for training, two for validation, and the remaining two were reserved for testing (same as in the split described in 6.5.4), resulting in a training set of 25 membranes and a validation set of eleven membranes.

GT segmentation masks were generated using the spheres approach, where spheres of radius 4 voxels were placed at the 3D GT positions. DeepFinder was trained using a patch size of 40 voxels for 100 epochs with 20 steps per epoch and a step size of 9 voxels for random shifts. After training, segmentation masks were predicted for the test tomograms, and protein positions were extracted using DeepFinder’s clustering module with a bandwidth parameter of 3 voxels.

All mentioned hyperparameters were optimized through grid search using the validation tomograms. To associate predicted protein positions with membranes, we considered only positions within a maximum distance of 4 voxels from the membrane surface.

#### CrYOLO

For CrYOLO^52^, we used the same data split as for DeepFinder, ensuring non-overlapping training and validation patches per tomogram. GT positions were converted into CrYOLO-compatible format by generating .cbox files containing bounding box coordinates. A box size of 10 voxels was chosen, with bounding boxes propagated across three slices above and below each GT position.

Hyperparameter tuning was performed on the validation dataset to determine the optimal settings, resulting in the following parameters: threshold = 0.05, tracing_min_length = 3, distance = 6, min_num_boxes = 5, and positive_weight = 50.

Membrane-position associations were performed following the same approach as for DeepFinder, considering only predicted positions within 4 voxels of the membrane surface.

#### MPicker / EPicker

For the comparison with MPicker^58^ and EPicker^117^, we first flattened our membranes using the MPicker GUI. This involved manually selecting several points per membrane on the luminal side of the segmentation to define a reference surface on the correct membrane side. MPicker’s built-in functionality was then used to flatten the membrane, generating a 2D image stack representation. Next, we utilized MPicker’s “Load raw coordinates” function to import a text file containing our GT positions for PSII and UK particles. MPicker transformed these 3D coordinates to their corresponding positions on the flattened membrane stack (see <u>Supp. Fig.</u> <u>S13</u>). Finally, for each membrane, we saved the flattened membrane stack as an .mrc file and the transformed GT positions as a .txt file.

To train EPicker, we needed pairs of 2D images (.mrc files) and their corresponding 2D coordinates (.thi files). These were prepared by extracting the 2D slice from the flattened membrane array that contained the most protein center positions. Additionally, we included the coordinates from the adjacent slices in both directions along the z-axis, as the center slice still retained enough protein density to confirm the presence of these proteins.

EPicker training was conducted using a learning rate of 0.0001, a batch size of 4, and 140 epochs, executed via the script `epicker_train.sh`. We trained separate models for each of our four training folds, matching the same folds used in the MemBrain-pick and MemBrain v1 analyses. After training, we predicted positions on the validation membranes and optimized the parameters `--thres` (score thresholding) and `--dist` (minimum distance between detected particles) using the Mpicker `epicker_batch.py` script. These parameters were fine-tuned separately for each data fold.

Once the optimal parameters were determined, we applied EPicker to our test set. MPicker then converted the predicted 2D coordinates back into 3D tomogram coordinates, enabling direct comparison with the GT positions. This allowed us to compute performance metrics and assess the accuracy of the MPicker-EPicker pipeline relative to our GT annotations.

---

**Supplementary Table 1.** Overview of data folds in ablation studies. Columns represent different datasets (Spinach, Chlamy, Community, Synthetic, DeePiCt), and rows indicate the number of tomogram patches allocated to each cross-validation fold. The “Total” row shows the cumulative number of patches per dataset.

| Fold \ Dataset | Spinach | Chlamy | Community | Synthetic | DeePiCt |
| --- | --- | --- | --- | --- | --- |
| Fold 1 | 10 | 7 | 5 | 8 | 4 |
| Fold 2 | 12 | 6 | 6 | 8 | 4 |
| Fold 3 | 17 | 7 | 5 | 8 | 4 |
| Fold 4 | 19 | 7 | 5 | 8 | 0 |
| Fold 5 | 11 | 6 | 6 | 8 | 3 |
| Total | 69 | 33 | 27 | 40 | 15 |

**Supplementary Figure S1.**
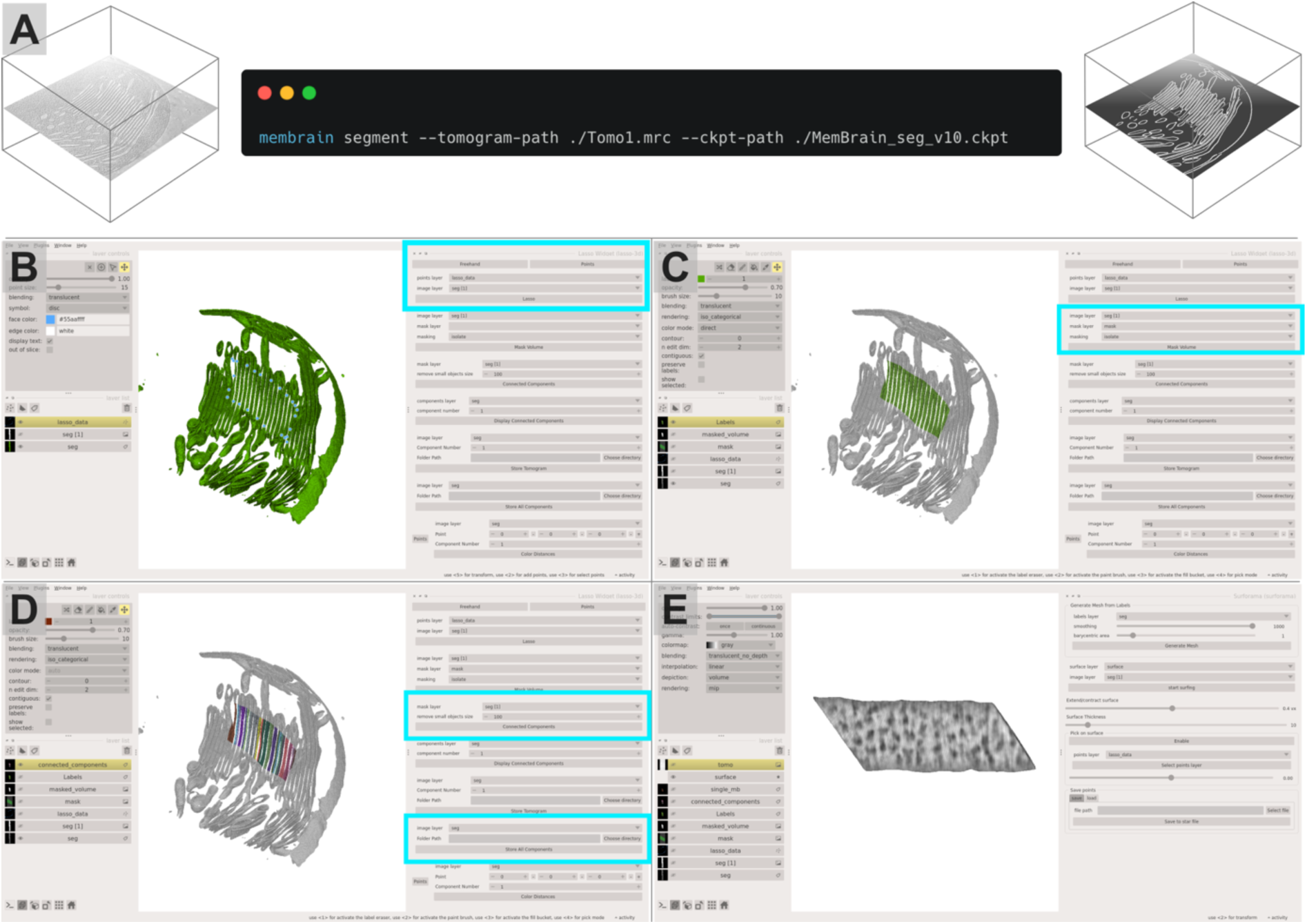
Napari plugins. **A:** MemBrain-seg can be used with a single command line `membrain segment` and outputs binary segmentations from a tomogram. **B-D:** MemBrain lasso plugin. **B**: Click and drag to draw a 3D lasso to select an area of interest. **C:** Either isolate or remove the selected area from the segmentation. **D**: Extract connected components of the remaining segmentation to find the membrane instances of interest, and save them out as .mrc files. **E**: Extracted membranes can be visualized in Surforama.

**Supplementary Figure S2.**
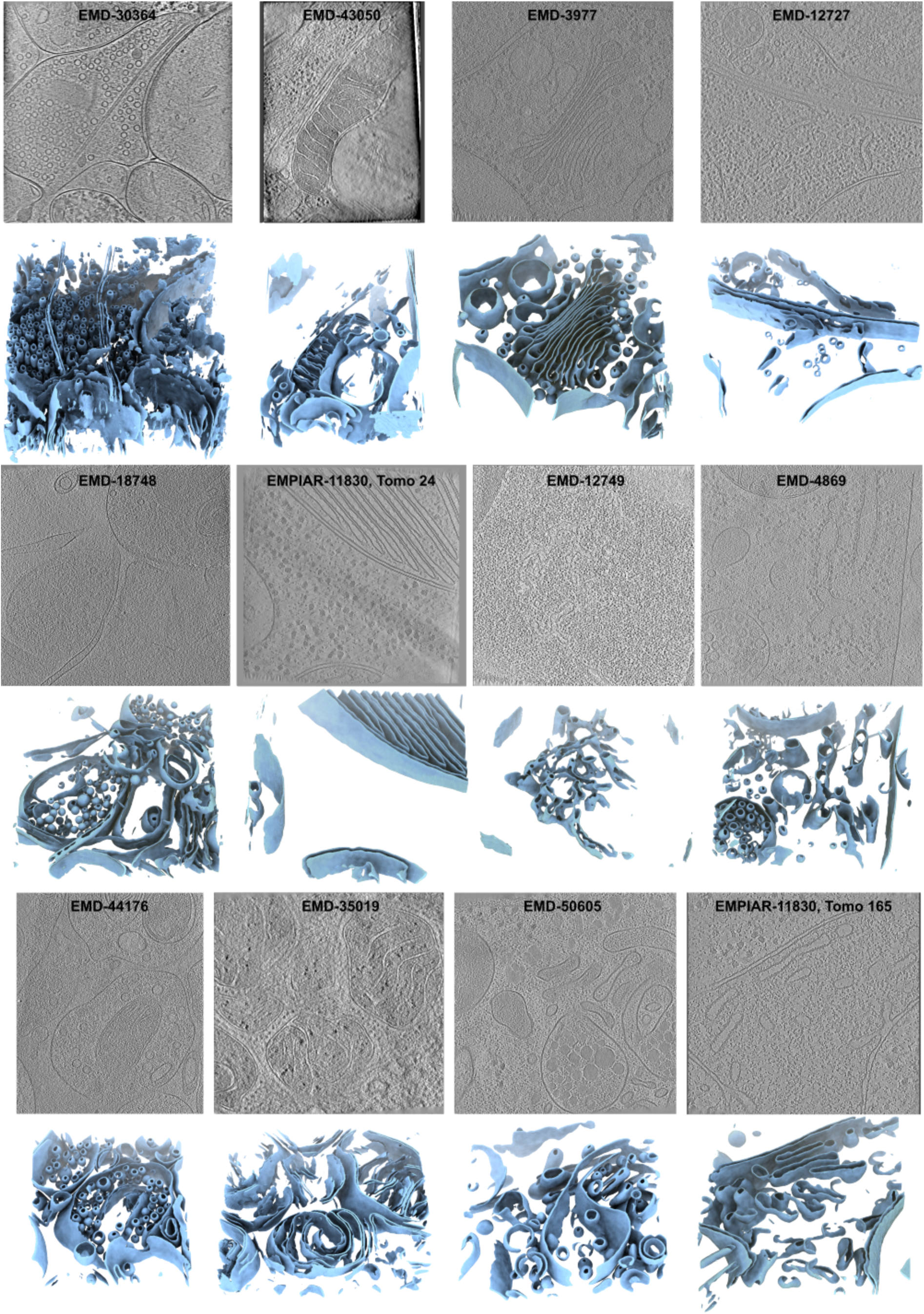
More examples of MemBrain-seg segmentations from different datasets and data sources. For each EMPIAR or EMDB dataset shown, top row: slice through tomogram, bottom row: corresponding MemBrain-seg prediction (light blue).

**Supplementary Figure S3.**
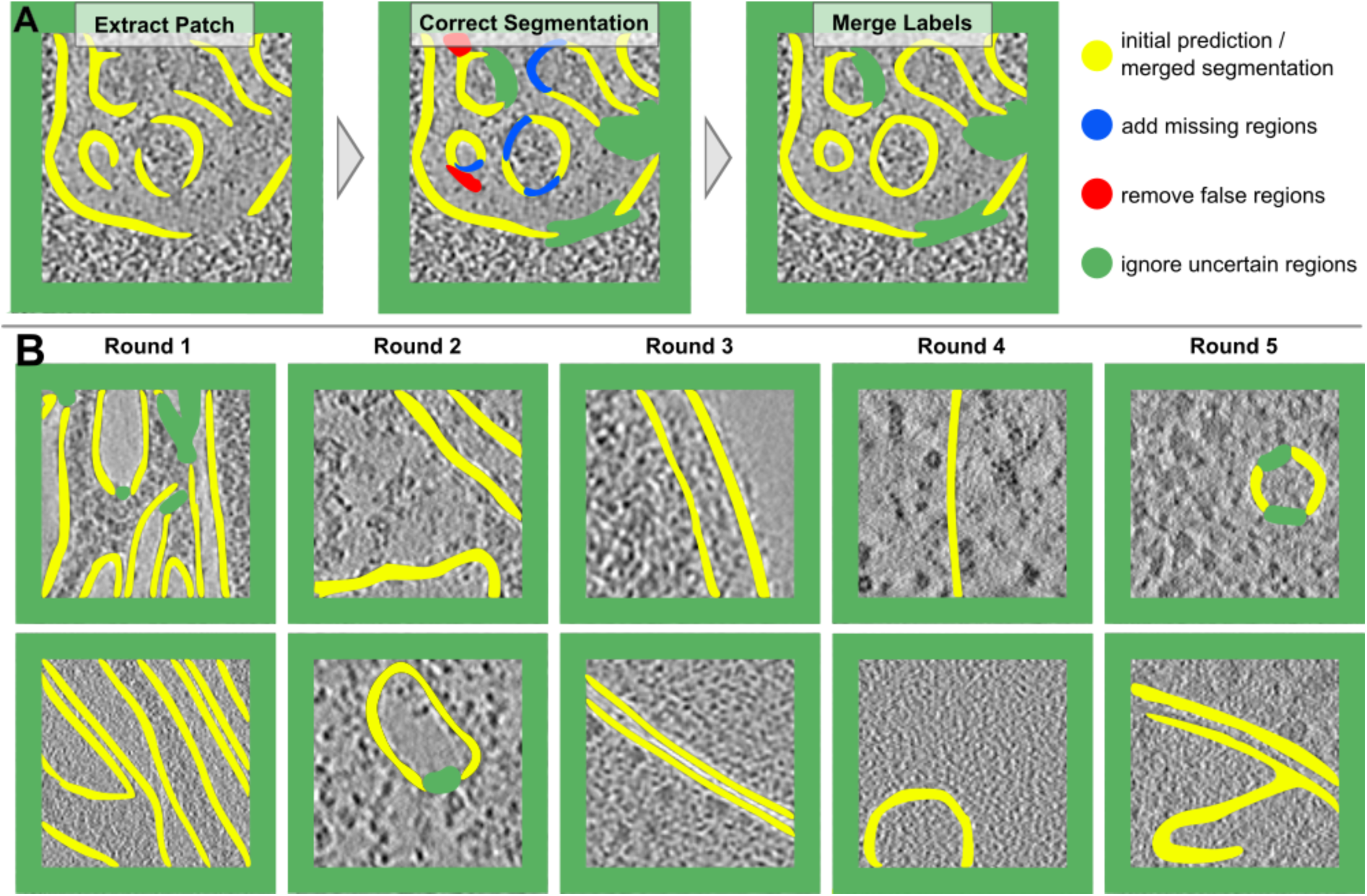
MemBrain-seg iterative annotation process. **A:** Correction of a single patch. An initial prediction of MemBrain-seg (left) is padded with ignore labels towards its edges to account for lack of context towards the edges. The correction (middle) includes added segmentations (blue), removal of false positive segments (red), and ignore labels in uncertain regions (green), where it’s not possible to perfectly delineate the membrane. The merged segmentation is visualized on the right. **B:** Example corrected patches from the datasets added in each iterative training round.

**Supplementary Figure S4.**
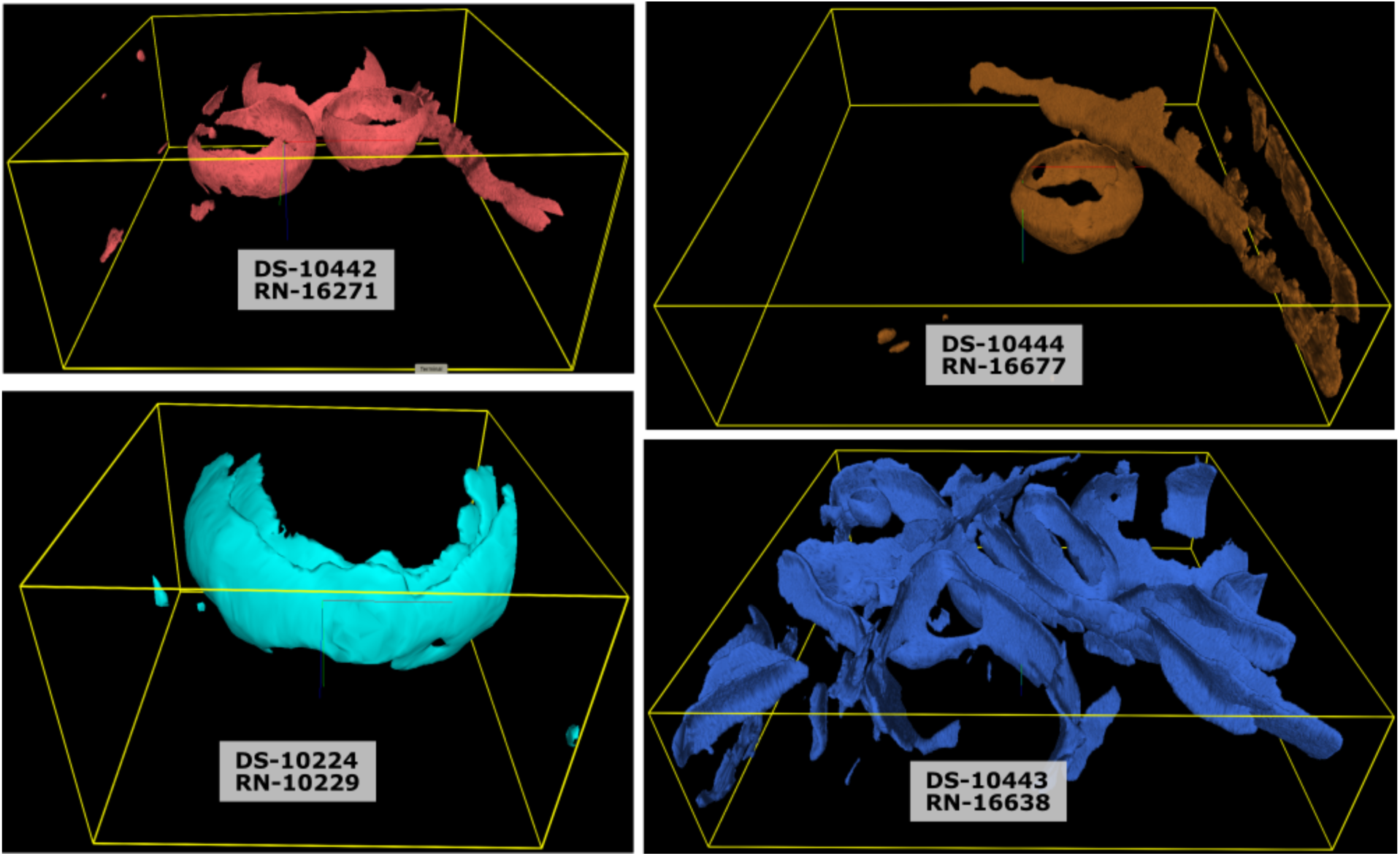
MemBrain-seg segmentations on the CZII CryoET Data Portal. All tomograms on the portal include MemBrain-seg segmentations, which can be viewed directly in the browser via Neuroglancer^118^. Shown are example segmentations from datasets DS-10442, DS-10444, DS-10223, and DS-10443.

**Supplementary Figure S5.**
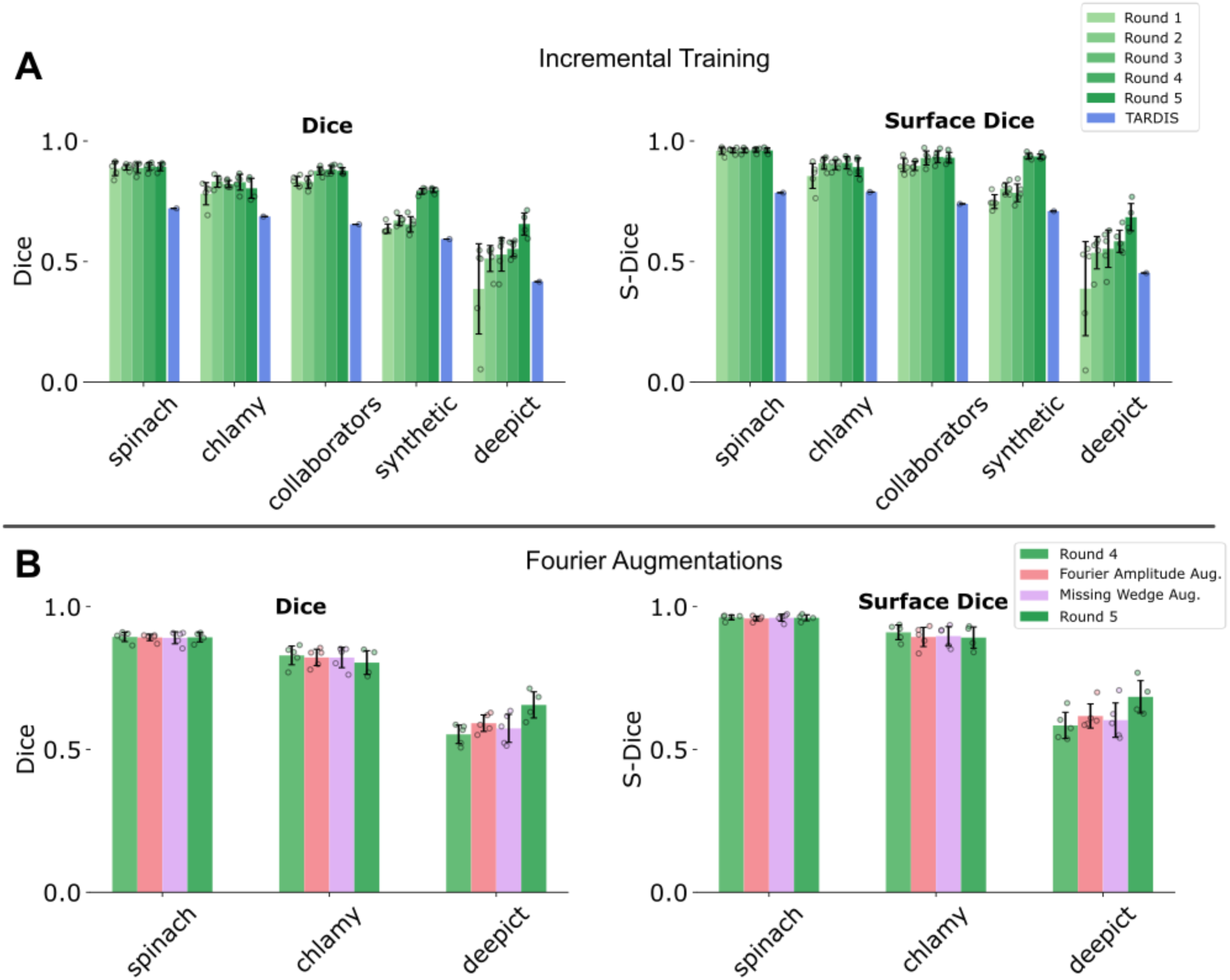
MemBrain-seg Evaluations. **A:** Evaluation of incremental training performance on different datasets with Dice and Surface-Dice scores, and comparison with TARDIS. MemBrain-seg models were trained with increasingly bigger training sets from rounds 1 to 5. **B:** Evaluation of the effect of our Fourier-based augmentations on generalization performance. Both models for Fourier Amplitude and Missing Wedge augmentations were trained with data from incremental training round 4, and compared to plain incremental training with rounds 4 and 5. All means, standard deviations and single points correspond to runs of the 5-fold cross-validation per experiment. TARDIS was evaluated only once on the entire dataset.

**Supplementary Figure S6.**
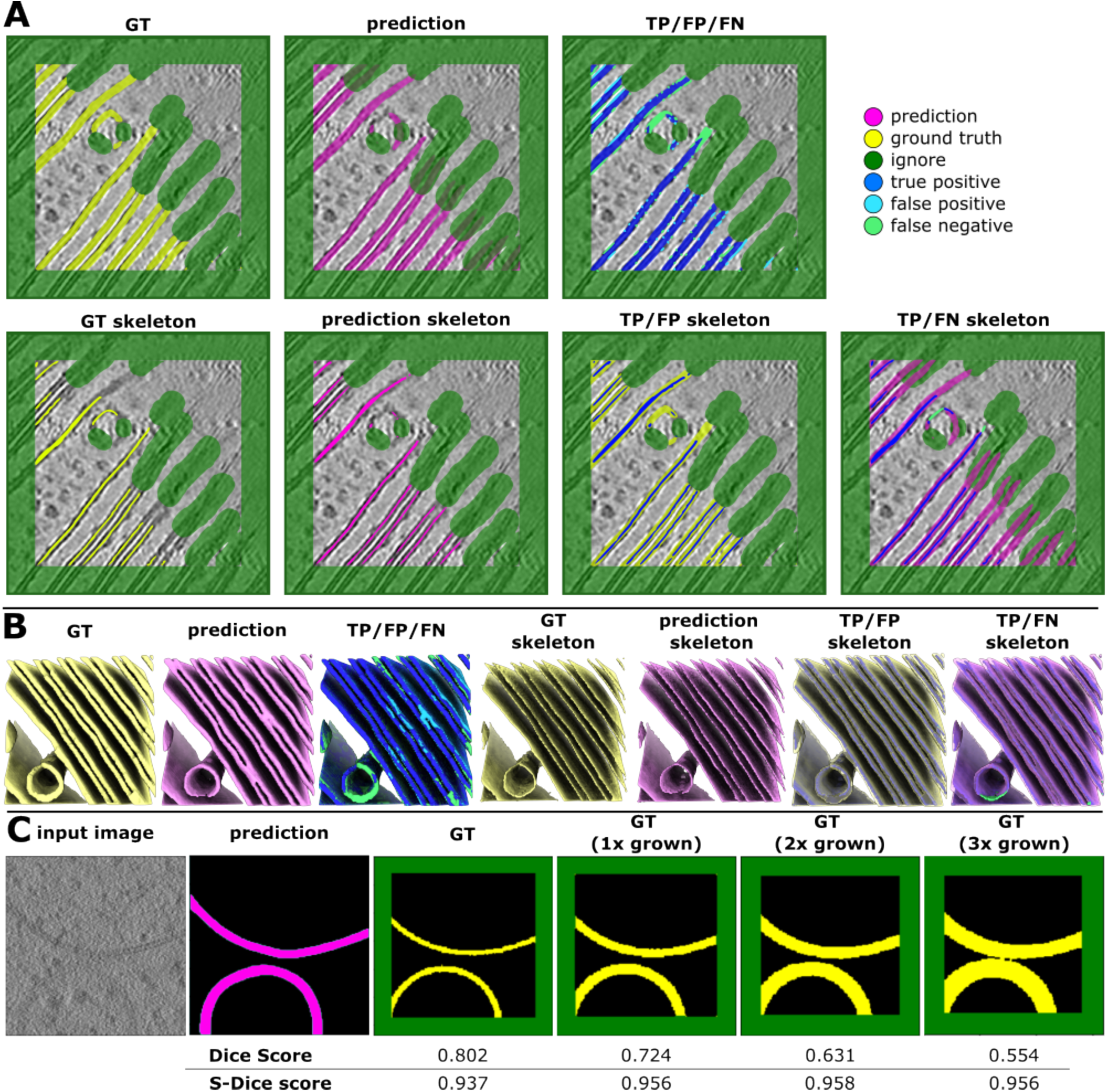
Surface-Dice. **A:** 2D slices of a corrected training patch. Comparison of *full* GT and prediction segmentation leads to false positive (FP) and false negative (FN) pixel for non-perfect agreement of segmentations, particularly visible at the segmentation edges. Bottom row: Comparing prediction skeletons with *full* GT segmentation (and vice versa) leads to bigger focus on membrane topology and gives false positives / negatives only in wrongly captured membranes. **B:** Same components as in A, but in 3D. **C:** Computation of Dice and Surface-Dice on a synthetic membrane patch: Both Dice score and Surface-Dice scores are computed by comparing a fixed predicted membrane segmentation with GT segmentations that is grown up to three times, leading to decreasing Dice scores, but consistent Surface-Dice scores.

**Supplementary Figure S7.**
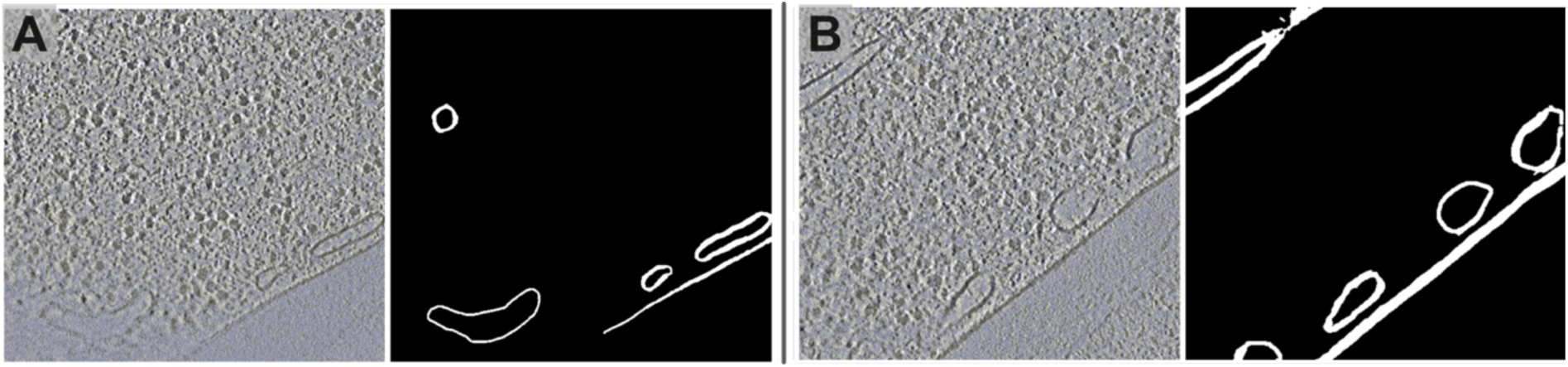
Differences in annotation thicknesses. Example segmentations provided in the DeePiCt dataset. Even though membrane topology is correctly captured, segmentation thickness varies among the different membrane instances.

**Supplementary Figure S8.**
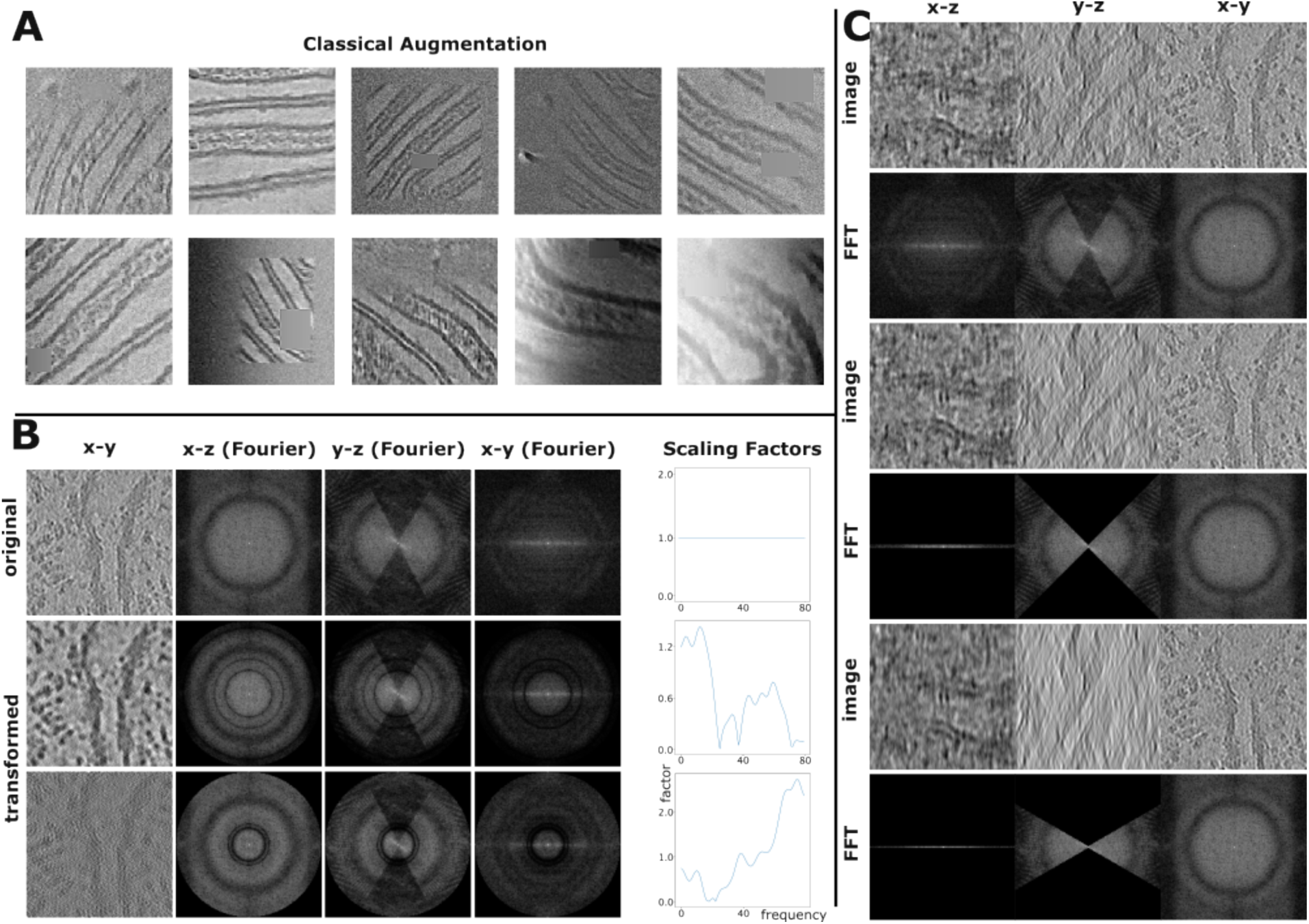
MemBrain-seg augmentations. **A:** Combinations of classical geometric and intensity transforms used in MemBrain-seg. **B:** Fourier Amplitude Augmentation: A patch and its Fourier transform is shown for the original patch (row “original”) and the same patch after drawing two random 1D plots (column “Scaling Factors”) and applying the corresponding rotational kernel. **C:** Missing Wedge Augmentation: A patch and its Fourier transform is shown for the original patch (rows 1 and 2), and for the same patch after applying an artificial missing wedge with different strengths (rows 3&4 and 5&6).

**Supplementary Figure S9.**
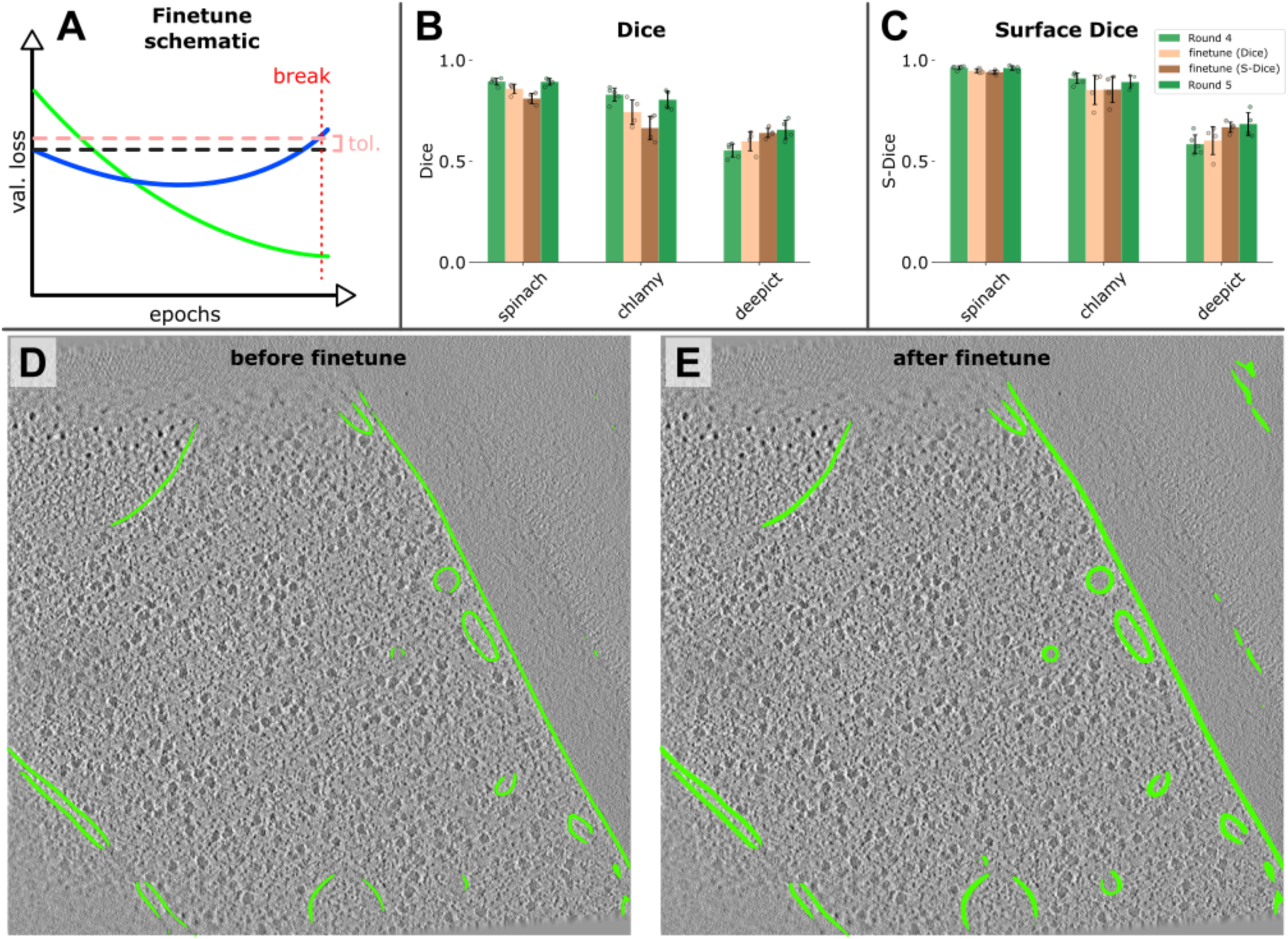
MemBrain-seg finetuning. **A:** Schematic of finetuning learning curves: Green training curve decreases while blue validation curve first decreases and then increases, indicating overfitting. We implement early stopping when validation performance deviates too far from original performance. **B:** Dice scores of incremental training rounds 4 and 5, compared to finetuning of a model trained using data from round 4 using only DeePiCt data. Finetuning models were trained with only Dice and BCE as loss function, as well as with added Surface-Dice. **C:** Surface-Dice of models described in B. **D:** Prediction on DeePiCt’s tomogram TS_0002 before finetuning (i.e. Round 4). **E:** Prediction on tomogram from D after finetuning with Surface-Dice.

**Supplementary Figure S10.**
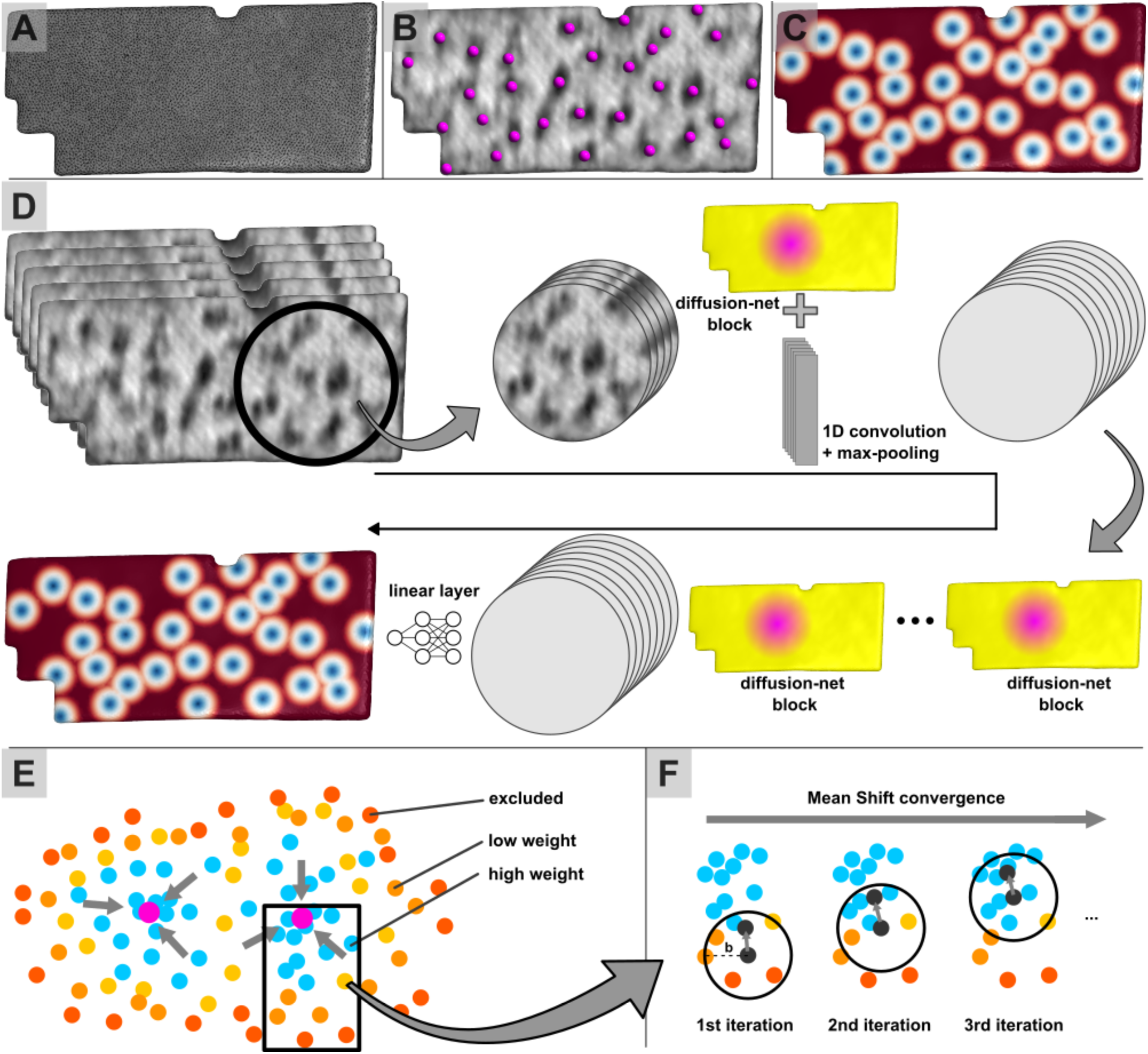
MemBrain-pick processing steps. **A:** Triangular mesh representation of a segmented membrane. **B:** Tomographic densities projected onto the mesh with GT particle positions (magenta). **C:** Training target: Geodesic distance map to the nearest particle center. **D:** MemBrain-pick architecture: Tomographic densities are projected onto the mesh from multiple distances from the membrane surface, creating *N* feature channels per vertex. The mesh is partitioned into overlapping, evenly sized regions to ensure shape consistency and prevent overfitting to global structures. Each partition is first processed by a 1D convolution along the channel dimension, followed by four DiffusionNet blocks. Finally, an MLP predicts the geodesic distance map to the nearest particle center. **E:** Score-guided mean shift clustering refines particle localization. **F:** Example iterations of a clustering seed (black) converging toward a predicted particle center.

**Supplementary Figure S11.**
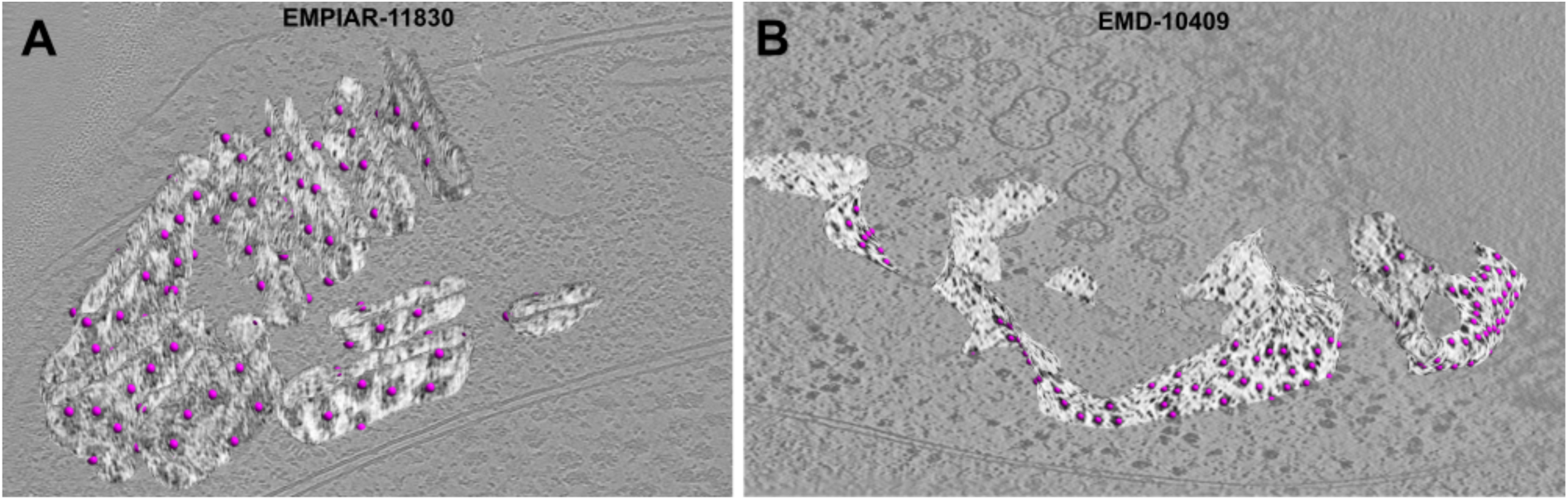
MemBrain-pick additional applications. **A:** MemBrain-pick predictions of respirasome particles on mitochondrial crista membranes in EMPIAR-11830^20,73^. **B:** MemBrain-pick predictions of ribosome positions on ER membranes in EMD-10409.

**Supplementary Figure S12.**
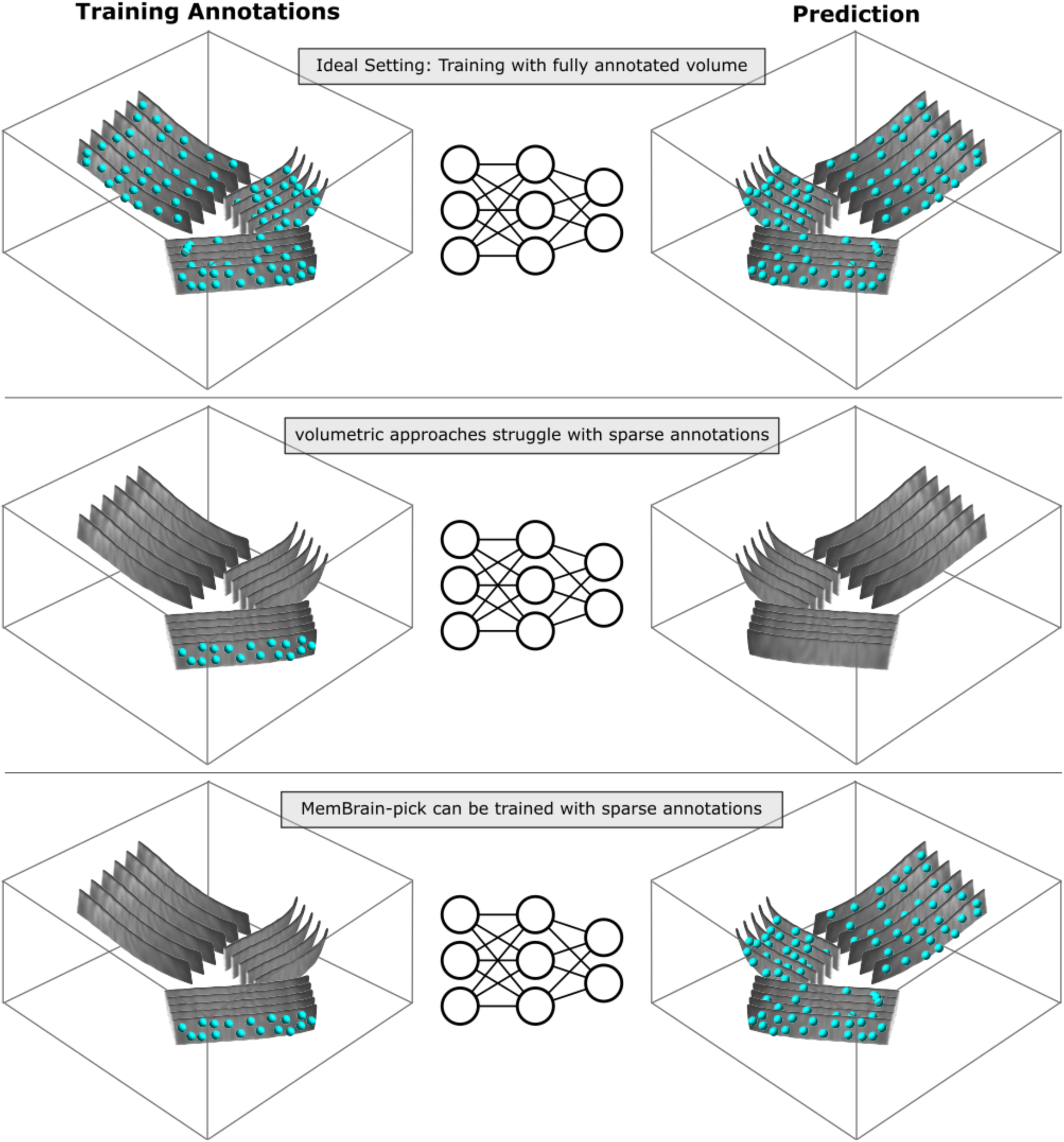
**Volumetric approaches vs. specialized approaches. Upper panel**: Volumetric approaches give good prediction results when trained with fully annotated tomogram regions. **Middle panel:** Volumetric approaches struggle to give good predictions when trained with sparsely labeled regions due to high amounts of false negative GT positions. **Bottom panel:** MemBrain pick and other membrane-specialized approaches require annotations only on a membrane-level instead of full-region annotations.

**Supplementary Figure S13.**
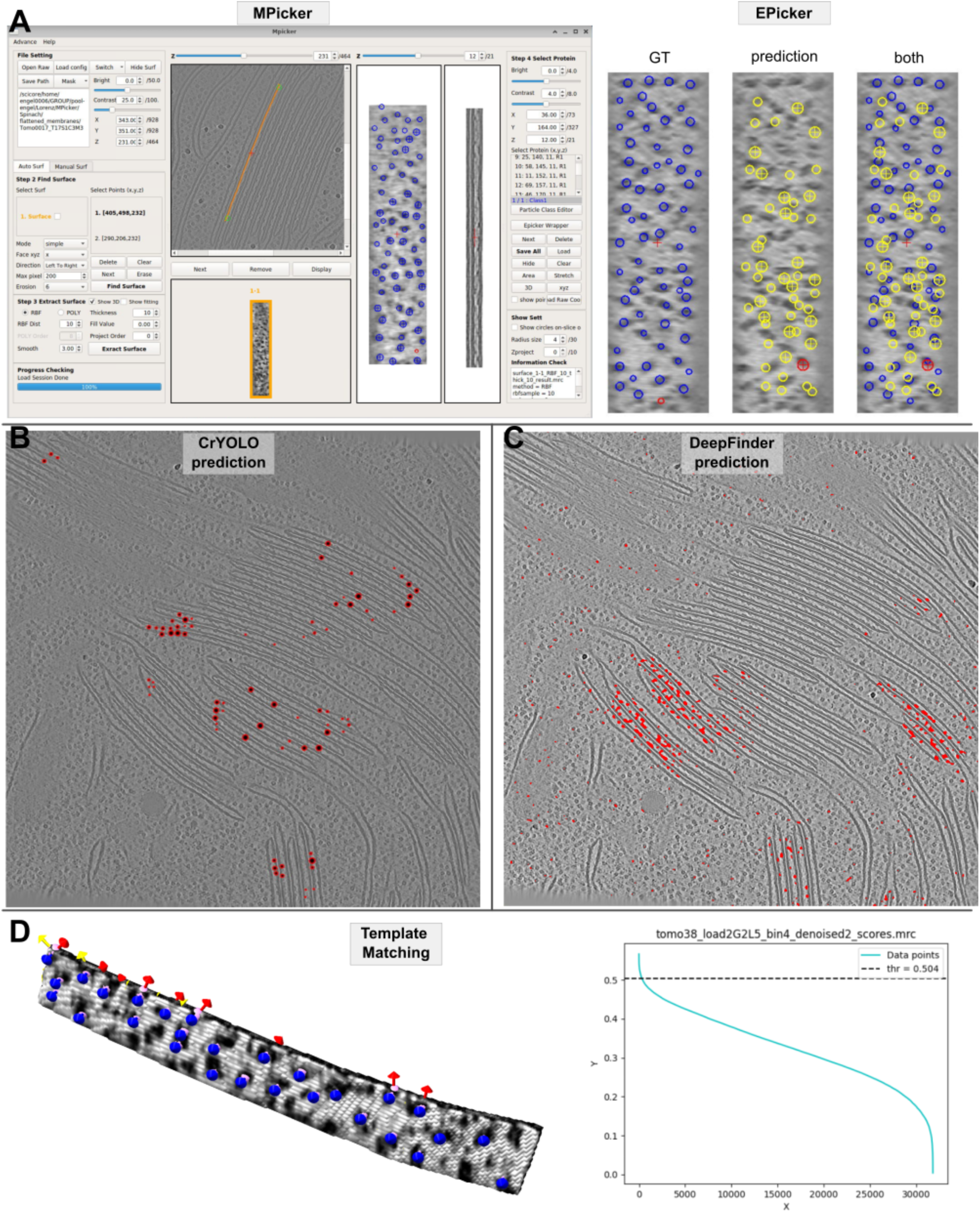
MemBrain-pick Compared Methods. **A:** Left: GUI window of MPicker visualizing a flattened training membrane from the Spinach dataset, together with our GT PSII positions mapped to the flattened 2D image. Right: Magnified view of GT positions on a Spinach test membrane, as well as positions predicted with EPicker. **B:** CrYOLO-predicted PSII positions on a spinach test tomogram, visualized in CrYOLO’s Napari plugin. **C:** DeepFinder predicted PSII segmentations on a Spinach test tomogram. **D:** PyTOM template matching (TM) results. Left: Example test membrane with detected positions (blue) mapped in. Right: sorted TM scores for this membrane with threshold used for extraction.

**Supplementary Fig. S14.**
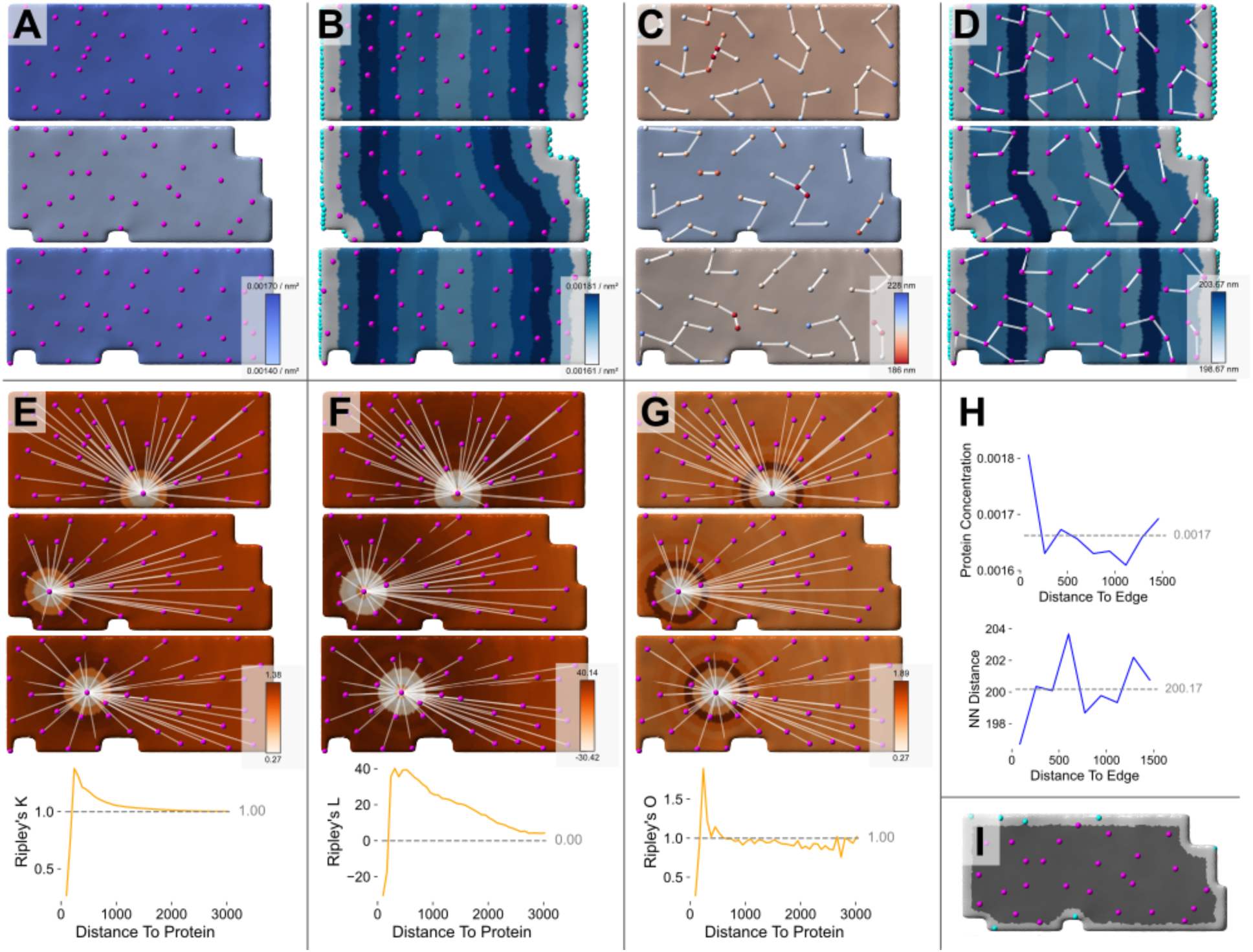
MemBrain-stats visualizations. **A:** Particle (magenta) concentrations for different membranes. Membrane surfaces are colored according to their concentrations. **B**: Particle concentrations with respect to distance to the membrane borders (cyan). Distances are divided in equally-spaced bins, and each bin is colored according to its concentration. Bin concentrations are computed from all membranes in one tomogram. **C:** Geodesic nearest neighbor distances: Membranes are colored according to their average nearest neighbor distances. Single particles are colored according to their respective nearest neighbor distances. White lines represent nearest neighbor connections. **D:** Membrane is divided into bins and colored according to average bin nearest neighbor distances, similar to B. **E:** Ripley’s K plot is visualized for a single starting protein. Membranes are colored according to their Ripley’s K value for the vertices’ geodesic distances to the starting proteins. **F:** Same as E for Ripley’s L. **G:** Same as E for Ripley’s O. **H:** Particle concentrations / Geodesic nearest neighbor distances per binned distances to the membrane edge, corresponding to the membranes shown in B and D. **I:** Segmentation edge exclusion: Particle positions close to the segmentation edge (cyan) can be excluded from some analyses.

**Supplementary Figure S15.**
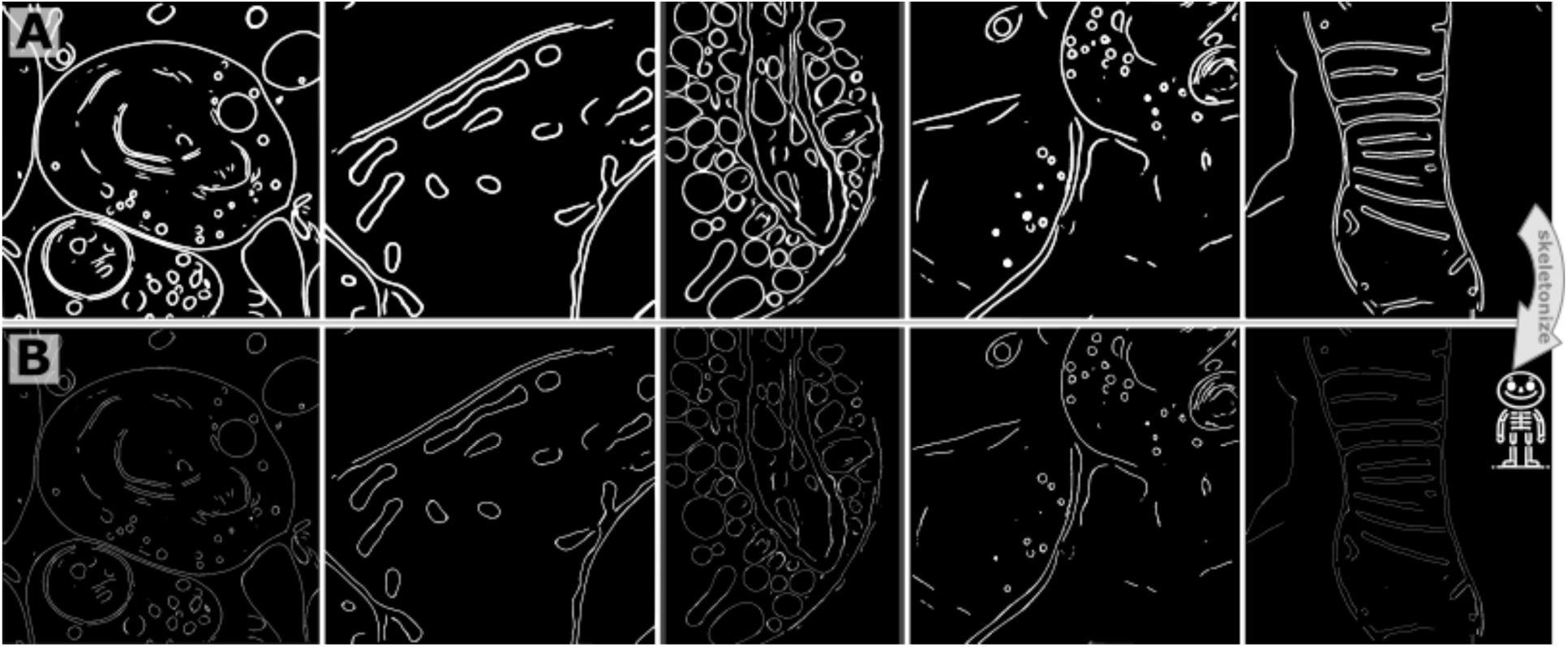
**MemBrain-seg skeletonization. Top row**: Example raw binary segmentation outputs from MemBrain-seg. **Bottom row**: The same segmentations skeletonized with the `membrain skeletonize` command.

**Supplementary Figure S16.**
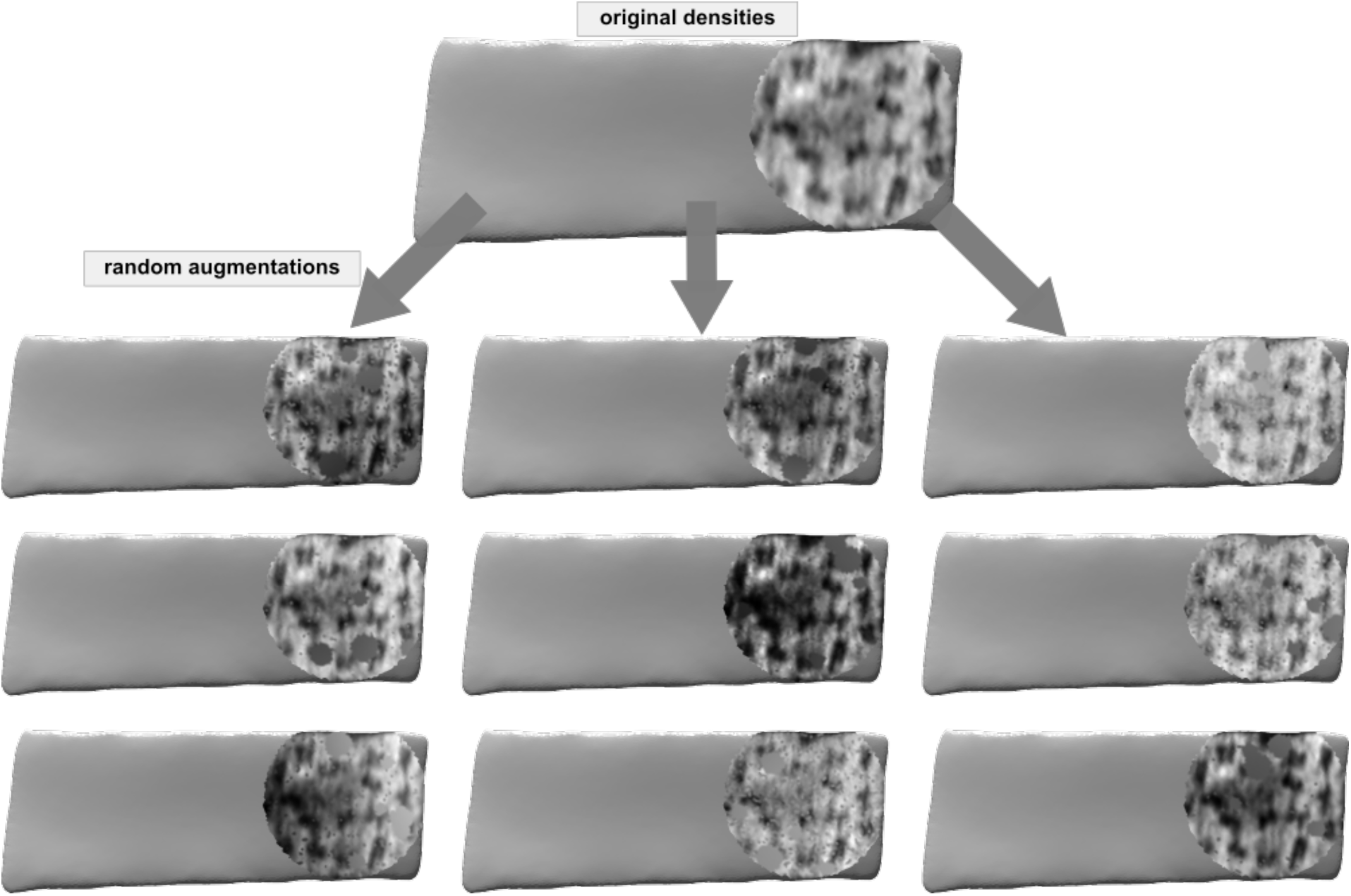
MemBrain-pick Augmentations. The top row shows a MemBrain-pick partition with one channel of original tomographic densities projected onto the surface. The rows below show examples of the same densities after random combinations of augmentations applied during training.

**Supplementary Figure S17.**
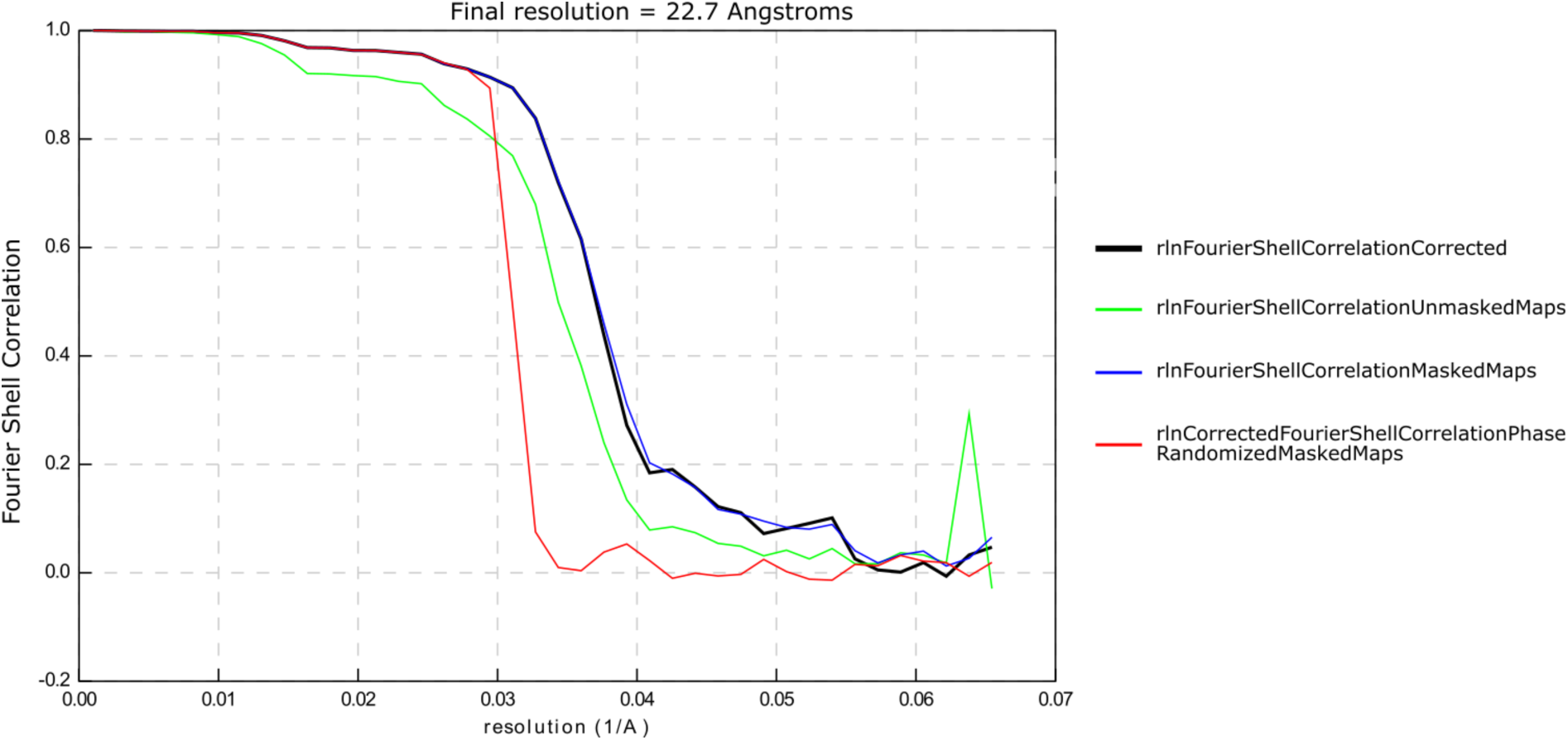
Fourier Shell Correlation (FSC) for the nuclear envelope-bound ribosome average shown in. Figure 5. The final resolution is 22.7 Å based on the 0.143 cutoff criterion after correcting for artificial correlations induced by the mask^119^. Shown are unmasked (green), soft-edged spherical mask (blue), corrected (black) and phase-randomized (red) FSC curves. The spike close to Nyquist frequency in the unmasked (green) curve is due to membrane signal extending all the way to the edge of the boxes since the half-map reconstructions not masked in STOPGAP.

